# Zfp263 is a transcriptional checkpoint of neutrophil development

**DOI:** 10.1101/2024.08.22.609132

**Authors:** Abhinandan Devaprasad, Zhichao Ai, Ananda Kishore Mukherjee, Áron Pánczél, Tariq Khoyratty, Ivan Ballesteros, Andrea Rubio-Ponce, Erinke van Grinsven, Lihui Wang, Julia Salafranca Gomez, Hayley L Eames, Andres Hidalgo, Attila Mócsai, Irina A Udalova

## Abstract

During development neutrophils undergo extensive changes in nuclear morphology, gene expression programmes, and acquire specific effector functions. However, molecular mechanisms controlling neutrophil development remain largely unknown. Here we systematically analysed changes in gene expression and chromatin landscape across stages of neutrophil differentiation, to construct a network of transcription factors, predicted to control neutrophil development and acquisition of functions. Zinc finger protein 263 (Zfp263) emerged at the apex of this network. Neutrophil maturation *ex vivo* and *in vivo* was blocked in cells lacking Zfp263 and accelerated if its levels were induced. Mechanistically, Zfp263 acts as a direct transcriptional repressor of the ERK1/2 pathway, that supports the proliferative capacity of neutrophil progenitors and limits their differentiation into mature cells. Blocking ERK1/2 signalling in the Zfp263 deficient cells using chemical inhibitors, rescues neutrophil differentiation. Other targets of Zfp263 include genes involved in neutrophil migration, cytokine and chemokine expression, reactive oxygen species and formation of neutrophil extracellular traps. Thus, our study outlines a framework for modulation of neutrophil development and / or function, aimed at neutrophil-mediated diseases.

## Introduction

Neutrophils are the primary responders to infection and inflammation due to their efficient production in the bone marrow (BM) and rapid mobilization to the blood and tissue (*1*, *2*). Upon reaching the inflammatory site, they execute crucial effector functions, including phagocytosis, the formation of neutrophil extracellular traps (NETS), generation of reactive oxygen species (ROS), and secretion of inflammatory factors and cytokines, to regulate immune responses (3–6).

Neutrophils continuous replenishment is ensured by a coordinated developmental process in the BM: from highly proliferative granulocyte-macrophage progenitors (GMP) (7), through distinct stages, including promyelocytic precursors, immature forms, banded neutrophils, to fully mature (segmented) non-proliferative neutrophils (8–10). These stages can be distinguished through morphological assessment and cell surface marker expression, with the markers like Ly6G, CD101, CD205, CD49d, CD81, etc., in mouse (9, 11–13) and CD66b, CD16, CD10, CD64, CD49d, etc in human (9, 14–16). Neutrophil maturation is predominantly governed by the granulocyte colony-stimulating factor (G-CSF) pathway (17). G-CSF, a hematopoietic growth factor, binds to its receptor, G-CSFR (CSF3R), and initiates a cascade of signalling events, ultimately activating mitogen-activated protein kinase (MAPK) cascades, with the ERK1/2 pathway activation serving as the primary effector (18, 19). Specifically, ERK1/2 acts as a gate-keeper of neutrophil differentiation: its suppression is needed to curb the proliferation of neutrophil precursors and direct them towards maturation (19).

Selected transcription factors (TFs), such as CCAAT/ enhancer-binding protein α (C/ebpα), Pu.1, growth factor independent protein 1 (Gfi1), Gata1, interferon regulatory factor 8 (Irf8), Runx1, Kruppel-like factor 6 (Klf6), and C/ebpε have emerged, often serendipitously, as important regulators shaping neutrophil development (20–26). Pu.1 and C/ebpα, were shown to guide neutrophil lineage commitment and early differentiation (24, 25); whereas C/ebpɛ and Gfi1 promoted neutrophil terminal differentiation (9, 26). However, despite the recent explosion of single-cell RNA analysis of neutrophil transcriptional states (27, 28), focussed elucidation of their molecular regulators has been missing. In this study, we systematically mapped the transcriptional and chromatin landscape of neutrophils during their development from GMP to mature segmented cells at high resolution and unravelled a network of transcription factors driving neutrophil development and associated functional programmes. Zinc finger protein Zfp263 emerged as a novel global controller of neutrophil maturation, acting as a checkpoint of the ERK1/2 signalling pathway and regulating the proliferation/differentiation switch.

## Results

### Chromatin changes underpin neutrophil maturation

To map out transcription regulators of neutrophil maturation and construct regulatory networks, we applied our recently developed neutrophil genomic pipeline (23) to a model system of immortalized HoxB8 myeloid progenitor cells, which allows for controlled differentiation into neutrophils with G-CSF (29). We have previously confirmed their similarity to primary neutrophils at different developmental stages in nuclear morphology, gene expression, and effector functions (14, 23). The cells were sampled at days 0, 1, 3, and 5 after stimulation with G-CSF, roughly matching GMP, metamyelocytes, banded and segmented neutrophil stages (14, 23), and subjected to ATAC-seq and RNA-seq analyses of chromatin and transcriptional changes respectively (**Figure 1A**).

**Figure 1:**
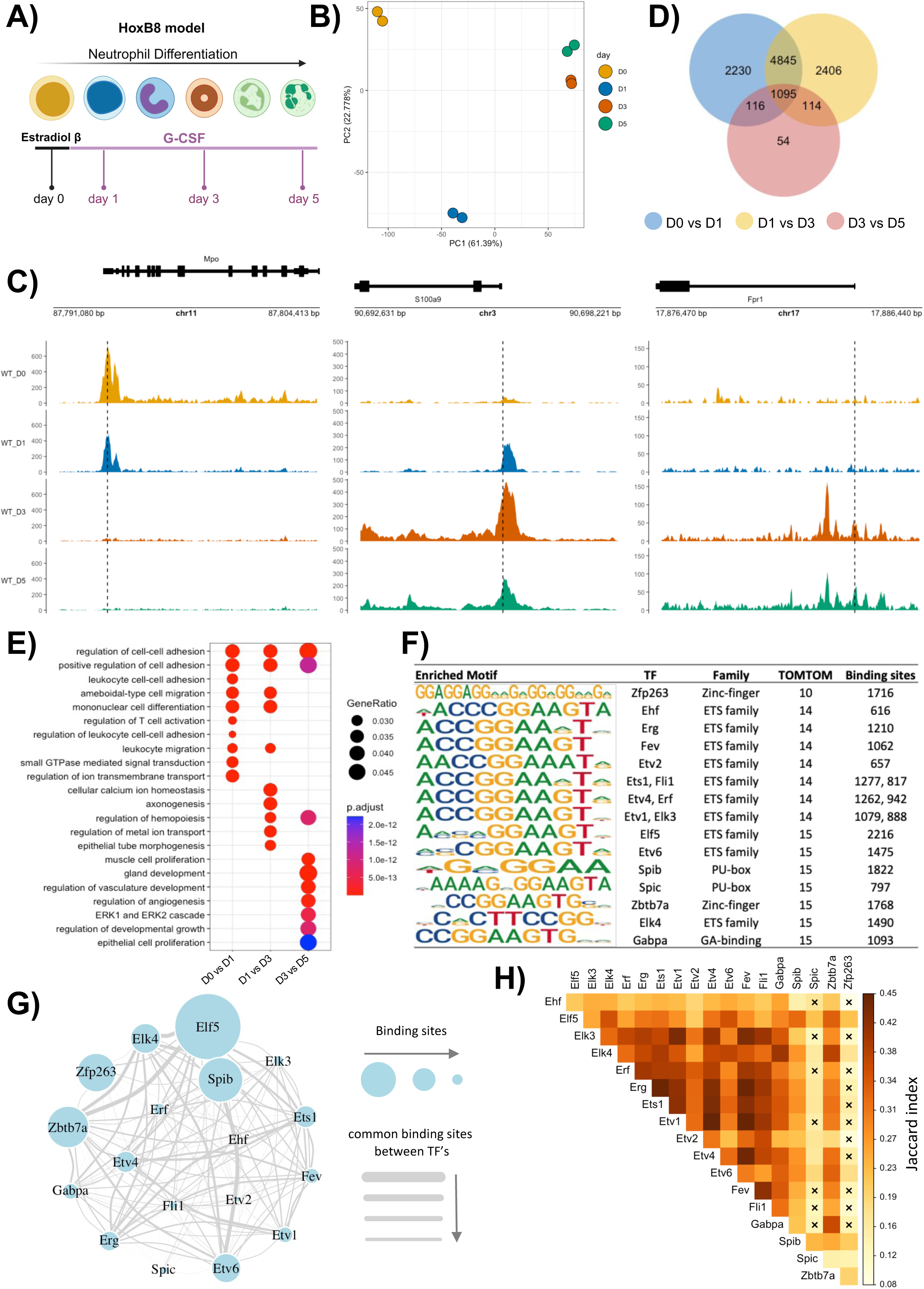
Neutrophils dynamically alter chromatin landscape during maturation. A. HoxB8 cells were differentiated and samples were collected at days 0, 1, 3, and 5. B. PCA analysis of 105394 DAPs (P_adj_ <0.05) derived from ATAC-seq analysis of HoxB8 differentiation at days 0, 1, 3, and 5 (n=4, replicates=2). C. Representative snapshots of gene accessibility based on normalized gene counts (FPKM). D. Venn diagram showing shared differential accessibility peaks (open and closed) across different days. E. Pathway analysis of differentially accessible genes between different time points comparison. F. Table describing top overlapping motifs from MEME-AME analysis with motif sequence of corresponding TF, TF-family, TOMTOM score, and number of binding sites. G. Regulon analysis of the key transcription factors (as shown in Figure 1F). Size of the nodes represents number of binding sites detected based on the MEME suite analysis in the DAPs proximal to TSS, across all time points. Thickness of the edges represents the common number of binding sites shared between the corresponding TFs. H. Heatmap representing the Jaccard’s similarity index for the binding sites shared between different TFs. The **x** represents TF pairs that a significantly different (Fisher’s exact test; p-value <0.05) in terms of the shared binding sites.

The PCA analysis of 105,394 differentially accessible peaks (DAPs; Padj < 0.05) derived from ATAC-seq analysis revealed that the chromatin of HoxB8 neutrophils has a distinct profile for each time point with days 0 and 1 displaying the biggest difference (**Figure 1B**). Hierarchical clustering of the DAPs identified five clusters of distinct patterns of chromatin behaviour, including the ones that either gained (clusters 1 and 5) or lost (clusters 3 and 4) accessibility with maturation, or had little/transient change (cluster 2) (**Supplementary Figure S1A**). For example, accessibility of MPO decreased with maturation. In turn, the accessibility of S100a9 increased initially with maturation and then decreased again. Whereas the Fpr1 locus opened with maturation (**Figure 1C**). We mapped DAPs to genes using GREAT (30) and performed reactome pathway enrichment analysis using ReactomePA from the clusterProfiler package (31, 32) on each cluster (**Supplementary Figure S1B**). The pathways enriched in cluster 1, with chromatin opening at days 1 and 3, comprised the RAC1 and RAC3 GTPase cycle. RAC GTPase play a crucial role in neutrophil adhesion, migration, and defence, thus important for neutrophil maturation (33–35). The pathways enriched in cluster 5, with chromatin opening at days 3 and 5, include toll-like receptor cascades, platelet activation, GPVI-mediated activation, and haemostasis. Cluster 3 and 4, characterised by chromatin closing over neutrophil maturation, were linked to pathways enriched for GTPases, ECM interactions, cell motility, WNT signalling, and phagocytosis. Cluster 2 was enriched for the haemostasis pathway (**Supplementary Figure S1B**).

Next, we mapped out pathways that were acquired or lost *en route* to maturity based on detected chromatin remodelling around the gene loci. 8286 genes differential between day 0 and day 1 were enriched in pathways associated with cell adhesion; 8460 genes differential between day 1 and day 3 were associated with leukocyte migration; and 1379 genes differential between day 3 and day 5 were associated with ERK1/2 cascade and regulation of cell proliferation (**Figure 1D, E**). DAPs in 1095 gene loci overlapped in all-time point comparisons (**Figure 1D**). These were inspected for known binding motif enrichment using MEME-AME (36) to ascertain putative transcription factors (**TFs**) with potential to control neutrophil maturation globally, across all stages of differentiation (see methods). 22 or 33 TF binding motifs were presented in the DAPs that consistently gained or lost accessibility over the days of neutrophil maturation; 18 TF binding motifs featured consistently in both types of DAPs (**Supplementary Figure S1C**). These belonged to the ETS, the zinc-finger, PU-box, and the GA-binding families. The most distinct motif based on the TOMTOM similarity index was Zfp263 binding motif (**Figure 1F and Supplementary Figure S1D**). We then performed regulon analysis, briefly, quantifying the number of DAPs proximal (± 2500 bp) to the transcription start site (**TSS**) which had binding for these 18 TFs. We found that Elf5, Spib, Zbtb7a, and Zfp263 bound to over 1500 DAPs (**Figure 1F, G**). Among these, Zfp263 and Spic shared fewer number of common binding sites with other TFs based on Jaccard similarity index and Fisher’s exact test (**Figure 1G, H**). A similar trend was observed when the regulon analysis was performed at different stages of neutrophil maturation with Elf5, Spib, Zbtb7a, and Zfp263 being the top TFs; Zfp263 and Spic consistently shared fewer number of common binding sites with other TFs across all time points comparisons (**Supplementary Figure S1E, F**). Between Zfp263 and Spic, Zfp263 had significantly higher number of binding sites across all time points (**Figure 1G, H and Supplementary Figure S1E, F**).

Thus, chromatin landscape undergoes extensive remodelling across all stages of neutrophil differentiation: from GMP to metamyelocytes to banded to segmented cells. These changes are linked to acquisition of specific neutrophil functions, including adhesion, migration etc and the loss of others, such as proliferation. A network of transcription factors has emerged as putative global regulators of these changes, with Zfp263 at its apex.

### Neutrophils show distinct gene expression profiles over maturation trajectory

Next, we mapped the transcriptional changes associated with neutrophil progression through the maturation trajectory by conducting RNA-Seq on the HoxB8 neutrophils at day 0, 1, 3, and 5 (**Figure 1A**). PCA of 8760 differentially expressed genes (**DEGs**) (LRT test, Padj < 0.05) clearly separated HoxB8 neutrophils from day 0, 1, 3, and 5 of *ex vivo* differentiation and PC1 showed a progression that correlated (R=0.91; p-value=0.0015) with the maturation trajectory (**Figure 2A**). The top or bottom 25 loadings from PC1 revealed genes, whose expression trends increased or decreased with maturation respectively (**Figure 2B**). For example, Fpr1 showed a gene expression trend that increased with maturation, whereas Mpo showed a trend that decreased with maturation - similar to its chromatin profile which gained accessibility with maturation (**Figure 1C, 2B**).

**Figure 2:**
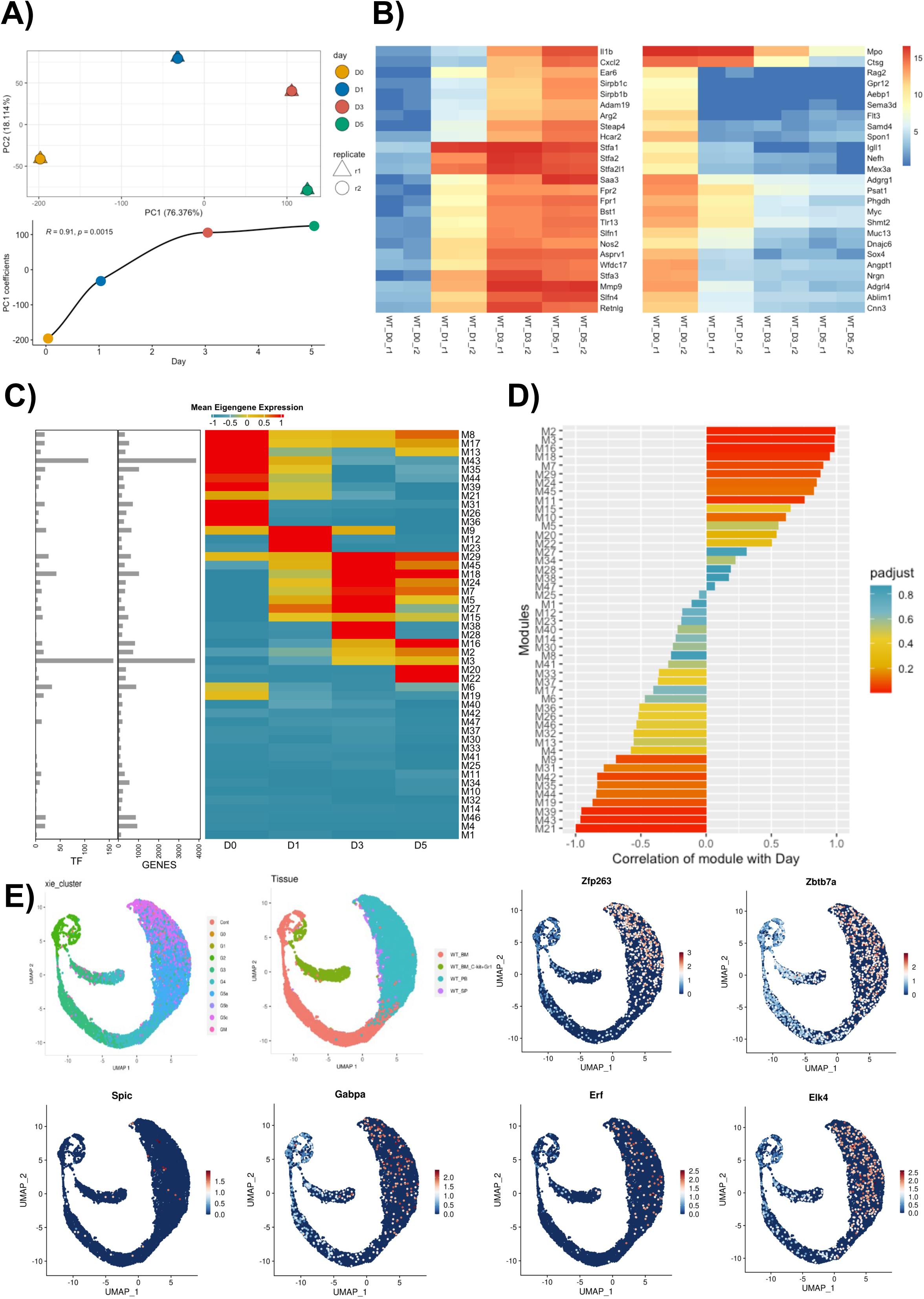
Neutrophil progression through the maturation trajectory is associated with distinct gene expression trends. A. Top panel represents PCA analysis of DEGs (P_adj_ <0.05) derived from RNA-seq analysis of HoxB8 differentiation at days 0, 1, 3, and 5 (n=4, replicates=2). DEGs were identified using the DESeq2 method. Bottom panel represents the coefficients of PC1 for all the samples. B. Gene expression heatmap of the top (left panel) and bottom (right panel) 25 loadings (genes) of the PC1. The colour of the heatmap represents gene expression based on variance-stabilizing transformation (VST). C. Heatmap showing mean eigen gene values of modules, side bar depicts number of genes and transcription factors present in each module. Module identification and eigen gene values were computed using WGCNA package. D. Pearson correlation between module mean eigen gene values of all modules vs. time. E. Expression of selected TFs in primary mouse neutrophils from bone marrow (BM), peripheral blood (PB), spleen (SP), with populations G0-G5 as defined by molecular signatures in the single-cell analysis of neutrophil heterogeneity (Xie et al. 2020).

To systematically identify genes and gene modules with time-specific gene expression trends, we performed weighted gene correlation network analysis (WGCNA), which revealed 47 gene modules (**Figure 2C**, see methods). Modules M2, M3, M16, M18, showed strong positive and modules M43, M21, M39 negative correlation with time (**Figure 2D**). The pathways enrichment analysis in modules positively correlated with time revealed programmes associated with mature neutrophil functions, such as inflammatory response, activation, response to infection, autophagy, and cytokine signalling (**Supplementary Figure S2A**). At the same time, the pathways enriched in modules negatively correlated with time related to cell biogenesis and included DNA replication, RNA processing, and ribosome assembly (**Supplementary Figure S2B**). Modules M2 and M3 displayed one of the strongest correlations with time (**Figure 2D**), and encompassed Zfp263, Zbtb7a, Spic, Gabpa, Etv1, Erf, and Elk4 TFs (**Supplementary Figure S2C**), which were highlighted by the chromatin landscape analysis (**Figure 1F**). Their expression progressively increased with time of HoxB8 neutrophil differentiation (**Supplementary Figure S2D**). When mapped onto the primary neutrophil populations from BM, blood, spleen, defined in previously published single cell profiling of neutrophil heterogeneity (27), expression of these TFs intensified with progression from cKit+Gr1+ proliferating progenitors, through BM populations maturation (G0-G4), reaching its maximum in most mature blood neutrophils G5a-c (**Figure 2E**).

*Zfp263* demonstrated one of the strongest differentials and levels of expression (**Figure 2E, Supplementary Figure S2D**). To better understand the regulatory potential of Zfp263 in neutrophil maturation, we constructed the Zfp263 expression correlation network using module M2 genes (**Supplementary Figure S2E**), that revealed the Zfp263 expression correlation with chemokine production, cytokine signalling pathway, cholesterol storage, regulation of Cxcl2 production, and ERK1/2 signalling cascade (**Supplementary Figure S2F**).

Thus, based on the accumulative evidence of the regulon (**Figure 1G, H**) and expression correlation network (**Supplementary Figure S2E**) analyses, we hypothesised that Zfp263 may control transcriptional programmes of neutrophil maturation.

### Zfp263 is critical for neutrophil maturation

To examine the role of Zfp263 in neutrophil maturation, we generated stable HoxB8 progenitor lines with loss of function (Zfp263KO) and gain of function (Zfp263AP) using CRISPR-Cas9 and validated the alteration of Zfp263 levels by western blot analysis (**Supplementary Figure S3A**). Following 5 days of *ex vivo* differentiation, wild type (WT) and Zfp263AP cells produced morphologically mature neutrophils, whereas Zfp263KO cells resembled a mix of largely immature and band neutrophils (**Supplementary Figure S3B**). Fluorescence-activated cell sorting (FACS) analysis confirmed that Zfp263KO cells generated a lower percentage of fully mature (Ly6G^hi^CD101^+^) neutrophils at day 5 (**Figure 3A**). Significantly, Zfp263AP cells reached the same level of maturity faster than the WT cells, at day 4 (**Figure 3A**), confirming a direct role for Zfp263 in G-CSF-driven neutrophil maturation *in vitro*.

**Figure 3:**
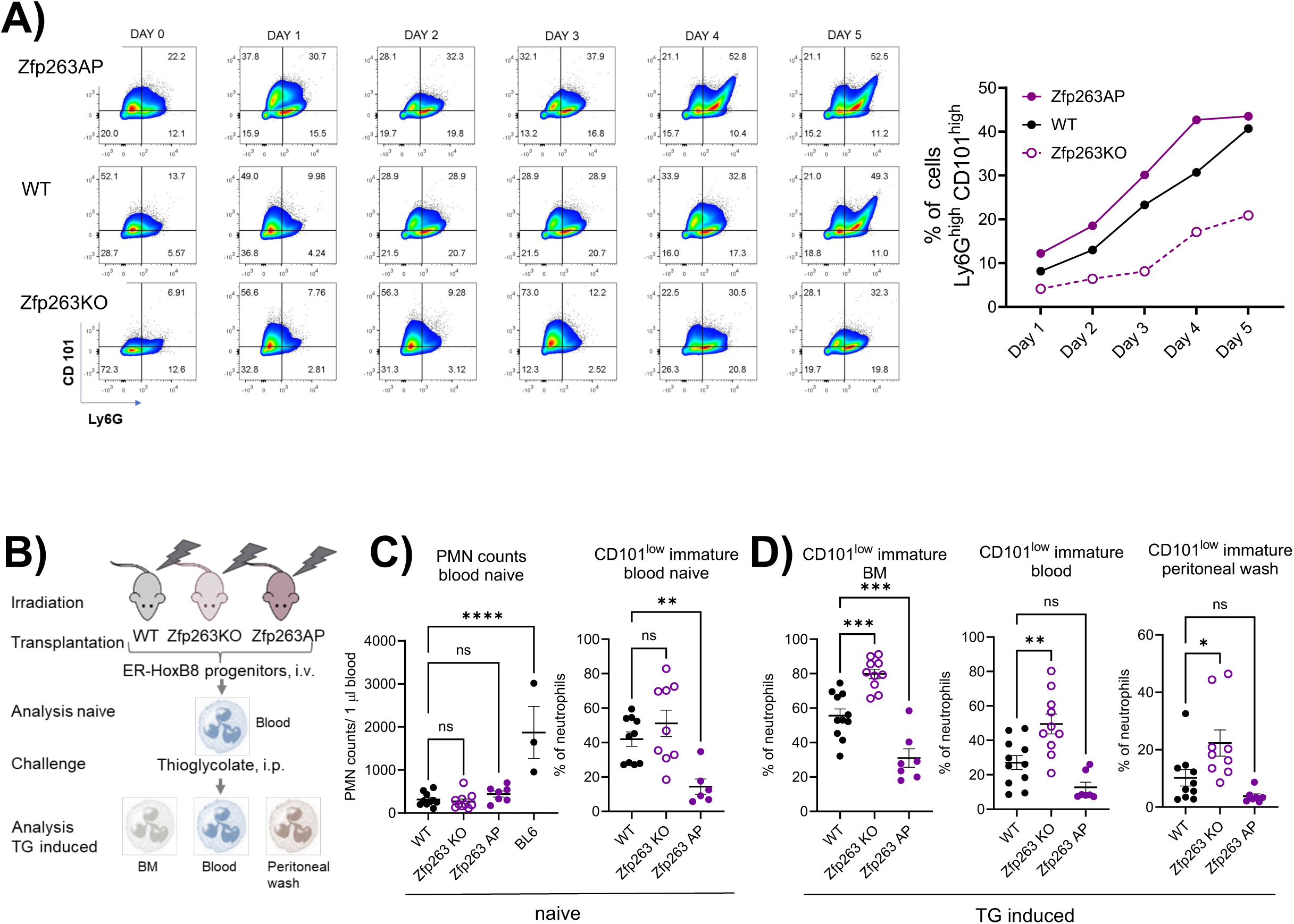
Zfp263 is critical for neutrophil maturation. A. Differentiation of HoxB8 cells (WT, Zfp263 AP and Zfp263 KO) was checked on Days 0,1,2,3,4 and 5 where Day 0 is the proliferating progenitor stage and Day 5 represents mostly mature neutrophils in WT. The expression of Neutrophil maturation cell surface markers Ly6G and CD101 were estimated by flow-cytometry. Ly6G^high^ CD101^high^ cells were gated to determine the % of mature neutrophils, as plotted in the right panel. B. Schematic depiction of the experimental setup: recipient mice were myeloablated by irradiation and transplanted with wild-type, Zfp263KO or Zfp263AP HoxB8 progenitors. Five days later, peripheral blood neutrophils were analysed by flow cytometry. Mice were then subjected to intraperitoneal (i.p.) thioglycolate (TG) injection and bone-marrow (BM), blood and peritoneum-infiltrating neutrophils were assayed again by flow cytometry. C. Comparison of absolute neutrophil counts of recipients of the indicated HoxB8 progenitors and intact wild-type mice (BL6) as well as proportion of CD101^low^ immature cells among neutrophils before thioglycolate challenge. D. Percentage of CD101^low^ immature neutrophils upon thioglycolate challenge in bone marrow, peripheral blood and at the peritoneal inflammatory site in recipients of the indicated HoxB8 progenitors. Statistical significance was calculated using one way ANOVA. Significant P value representation is as follows: * <=0.05, ** <=0.01, *** <=0.001, **** <=0.0001.

To further explore the potential of Zfp263 to drive neutrophil maturation *in vivo*, we have intravenously injected WT, Zfp263KO or Zfp263AP HoxB8 progenitors into lethally irradiated recipients and compared the ability of donor cells to engraft and develop into mature neutrophils (**Figure 3B**) (37). We detected similar levels of donor-derived neutrophil populations in the blood of recipient mice on day 5 post irradiation (**Figure 3C**). Consistent with the previously observed partial rescue of neutropenia by HoxB8 engraftment (37), neutrophil counts in chimeric mice were ∼2 times lower than in non-irradiated mice (**Figure 3C**). Flow cytometry confirmed a significant right-shift towards more mature neutrophils in the blood of mice that received Zfp263AP HoxB8 progenitors (**Figure 3C**). On the contrary, a trend towards increased frequency of immature neutrophils was observed in the Zfp263KO chimera mice (**Figure 3C**). Importantly, when the same mice were subjected to thioglycolate-induced sterile peritonitis, we registered a dramatic left-shift in neutrophil maturity in the Zfp263KO chimeric mice throughout the compartments: BM, blood, and the peritoneal wash (**Figure 3D**). In all analysed *in vivo* compartments, Zfp263AP neutrophils appeared to be more mature (**Figure 3D**).

Together, Zfp263 is functionally linked to neutrophil maturation: its depletion delays neutrophil differentiation *in vitro* and *in vivo*, while its overexpression expedites it.

### Zfp263 controls neutrophil immune responses and effector functions

To unravel the molecular events controlled by Zfp263 across the stages of neutrophil maturation, we performed ATAC-seq and RNA-seq on the Zfp263KO HoxB8 cells on day 1, 3 and 5 (**Figure 4A**). The comparative PCA analysis of the global ATAC-seq and RNA-seq profiles revealed that transcriptional and chromatin profiles of Zfp263KO cells clustered together with the WT cells sampled at earlier time points (**Figure 4B, C**), supporting the delayed maturation profile of the Zfp263KO neutrophils.

**Figure 4:**
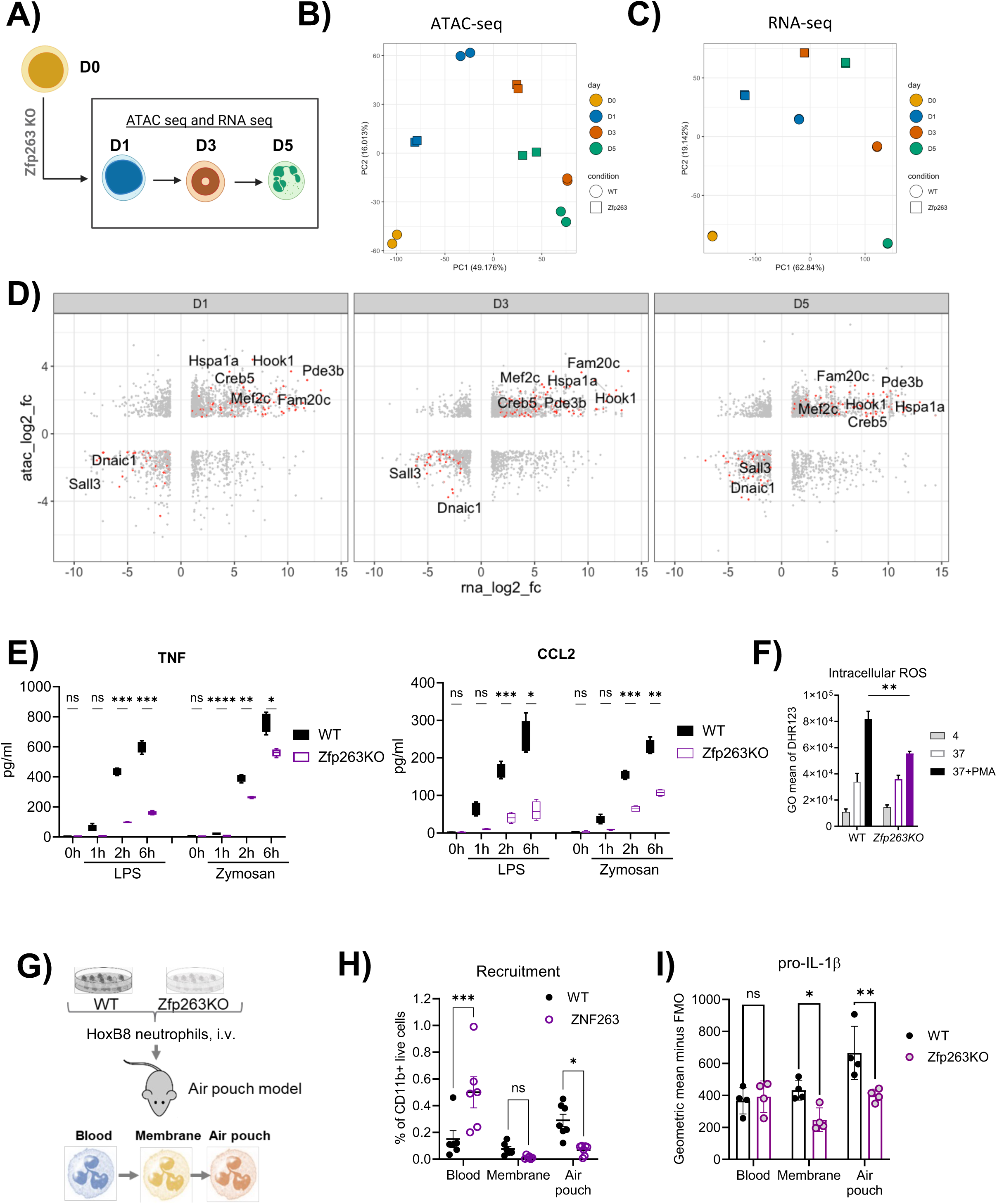
Zfp263 deficiency impacts neutrophil immune responses and recruitment to inflamed tissue. A. Experimental design for performing ATAC-Seq and RNA-Seq of WT and Zfp263 knock-out (KO) HoxB8 neutrophils. Samples (replicates=2) were collected at days 1, 3, and 5 for Zfp263 KO. B. PCA analysis of all DAPs (P_adj_ <0.05) derived from the ATAC-seq analysis of WT and Zfp263KO. C. PCA analysis of all DEGs (P_adj_ <0.05) derived from the RNA-seq analysis of WT and Zfp263KO. D. Combined differential analysis of chromatin and expression of WT vs. Zfp263. Orange dots indicate genes that are positively correlated between chromatin and expression, genes that have absolute fold change (log2) > 2 in both platforms have been labelled. E. Secretion of TNF and CCL2 in WT and Zfp263 KO HoxB8 cells with LPS and Zymosan stimulation assessed by ELISA. Statistical comparison was made by two-way ANOVA, *P<0.05, **<0.01. Data are shown as means and SD from four independent experiments. F. PMA induced intracellular ROS production by HoxB8 neutrophils were assessed by flow cytometry with dihydrorhodamine 123 (DHR123), and then analysed by normalising fluores-cence signals of stained cells to unstained cells. Wild-type and Zfp263-deficient HoxB8 myeloid progenitors were differentiated in medium supplemented with G-CSF to induce differentiation for 5 days, and it is followed by the overnight incubation with PMA to stimulate intra-cellular ROS generation. Comparison was made between WT and Zfp263KO HoxB8 neutrophils. Statistical comparison was made by one-way ANOVA, *P<0.05, **<0.01. Data are shown as means and SD from three independent experiments. G. CellTracker Far Red–labeled WT and CellTracker carboxyfluoresceinsuccinimidyl ester (CFSE)–labelled Zfp263KO HoxB8 neutrophils (Day 5 of differentiation) were intravenously injected in a 1:1 ratio. Following this, mice (N=6) subjected to air pouch formation were challenged with zymosan for 4 hours as previously reported (see methods). H. Labelled HoxB8 WT neutrophils from blood, air pouch membrane and air pouch exudate in mice (subjected to the air pouch model and zymosan challenge for 4 hours) were estimated by flow-cytometry and plotted as a percentage of total CD 11b + cells. Statistical significance was calculated by Mann-Whitney U test; significant P value representation is as follows: * <=0.05, ** <=0.01, *** <=0.001, **** <=0.0001. I. Expression of pro-IL1β, as measured by flow cytometry, in wild-type and Zfp263KO neutrophils recovered from blood, air pouch membrane and air pouch exudate of mice subjected to the air pouch model and zymosan challenge. All results are shown as means and SD derived from at least three mice within one experiment. Statistical comparison was made by one-way ANOVA, *P<0.05, **<0.01, ***<0.001, ****P<0.0001.

The correlative analysis on the ATAC-seq and RNA-seq Zfp263KO and WT datasets (**Figure 4D**) revealed 1062 differentially expressed genes with differential chromatin accessibility at day 1, 1258 at day 3 and 1157 at day 5 of *ex vivo* differentiation (**Supplementary Figure S4A**). Among these, 222 genes displayed differential gene expression and accessibility across all time points (**Supplementary Figure S4A**). 132 genes were positively correlated with expression and accessibility (represented by orange dots in **Figure 4D**), including Creb5, Dnaic1, Fam20c, Hook1, Hspa1a, Mef2c, Pde3b with an absolute log2 fold change > 2. The specific roles of these genes are not well characterized in the context of neutrophil maturation or function, hence further exploration is warranted. The GO pathway analysis of the differentially expressed genes across all time points (as shown in **Supplementary Figure S4A**) exposed an enrichment of pathways associated with signal transduction, specifically the ERK1/2 cascade and small GTPase signalling, as well as classical features of neutrophils related to the regulation of inflammatory response, adhesion, chemotaxis, and migration (**Supplementary Figure S4B**). Indeed, Zfp263KO neutrophils secreted less inflammatory cytokines and chemokines in response to LPS or zymosan stimulation, exemplified by TNF and CCL2 (**Figure 4E**). They also produced less intracellular and extracellular reactive oxygen species (ROS) in response to stimulation with PMA (**Figure 4F**) or fMLP **(Supplementary Figure S4C**), respectively. Neutrophils lacking Zfp263 were also deficient in NET formation, while activation of Zfp263 expression led to a slight elevation in NET production (**Supplementary Figure S4D**).

To validate the predicted impact of Zfp263 on neutrophil chemotaxis and migration, we utilised the previously described *in vivo* comparative quantification of neutrophil infiltration into the tissue lining and air pouch cavity in response to the zymosan (23). An equal mix of differently labelled WT and Zfp263KO HoxB8 neutrophils were injected intravenously prior to zymosan injection into the air pouch cavity (**Figure 4G**). No overall difference in cell survival, measured as Annexin V staining for apoptotic rate, was observed between the WT and Zfp263KO neutrophils (**Supplementary Figure S4E**). 4h post zymosan injection, we observed reduced infiltration of Zfp263KO neutrophils into the tissue lining and particularly into the air pouch cavity, while more Zfp263KO than WT neutrophils was detected in the circulation (**Figure 4H**), confirming the role of Zfp263 in promoting neutrophil trans-migration. Zfp263KO neutrophils in the membrane and exudate also produced less pro-IL-1β than WT neutrophils (**Figure 4I**), corroborating on their limited ability to produce inflammatory cytokines (**Figure 4E**).

In summary, Zfp263 deficiency impacts core neutrophil immune and effector functions, such as (trans)migration, cytokine, ROS and NET production. We speculated that Zfp263 controls neutrophil morphological and functional maturation via targeting key regulators of the corresponding transcriptional programmes.

### Zfp263 “target gene” network encompasses key regulators of cell differentiation

Considering a wide-reaching regulatory potential of Zfp263, including indirect regulation via networks, we wanted to distil direct targets of Zfp263. Thus, we scanned the DAPs in the vicinity of the **TSS** of all differentially expressed /remodelled genes (**Supplementary Figure S4A**) for the presence of the Zfp263 binding motif using the MEME suite analysis. 192 putative direct “target genes” were identified and used to generate the Zfp263 direct regulatory network at each time point in the maturation trajectory (**Figure 5A**, see methods). The colour of the edge represents the time point at which Zfp263 was predicted to bind to the gene and was differentially expressed between WT and Zfp263KO (**Figure 5A, B**). The shape of the nodes depicts TFs (diamond) or genes (circle) (**Figure 5A, C**). The total number of nodes in the Zfp263 network was significantly higher on day 1, compared to day 3 and day 5. 22 predicted Zfp263 “target genes” were TFs, including key transcriptional regulator of neutrophil maturation, such as Gfi1 (9, 26) and other recently highlighted maturation-related TFs, e.g. Klf6 (23) (**Figure 5D**).

**Figure 5:**
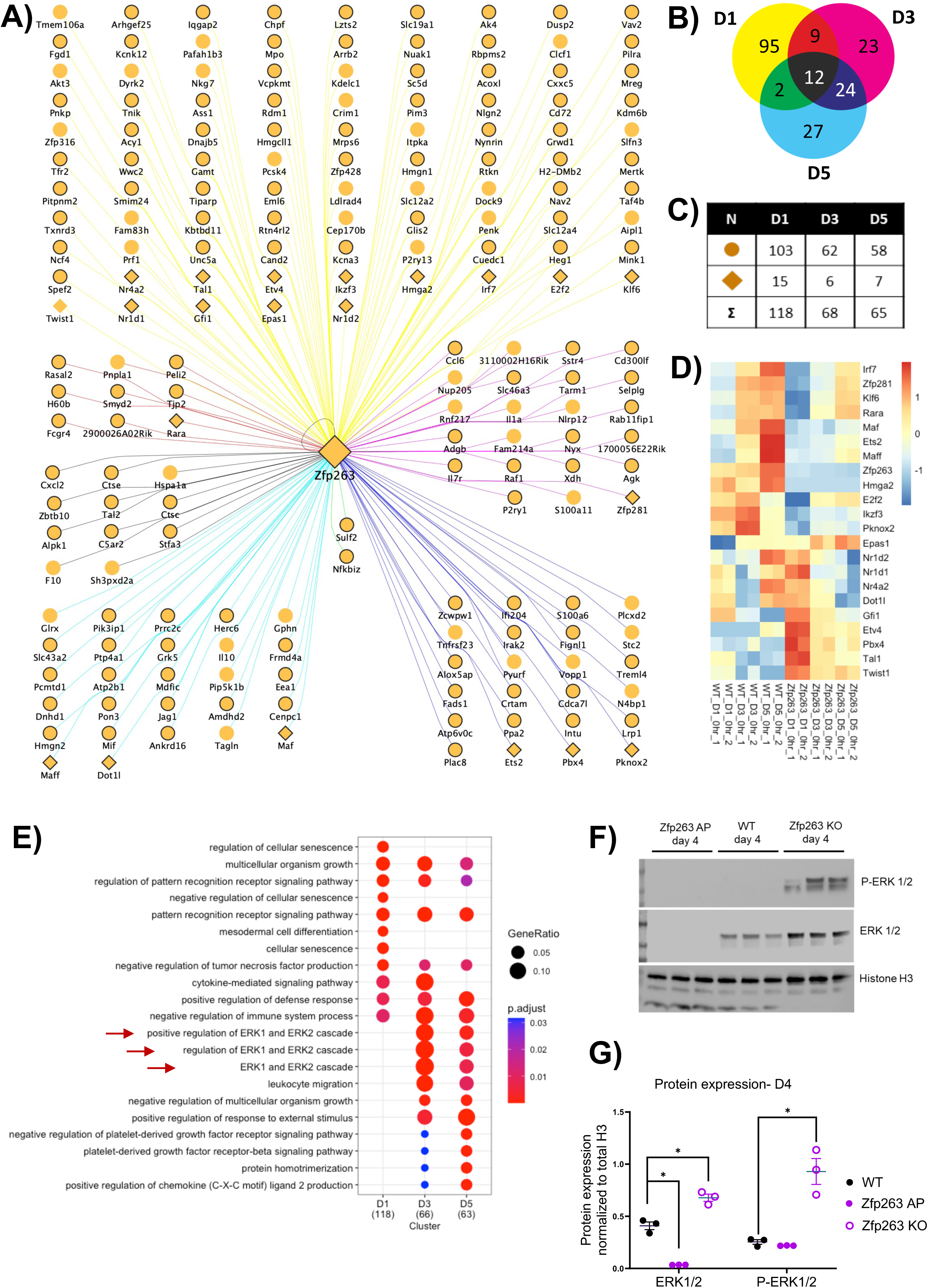
Zfp263 “target gene” network links Zfp263 and the ERK1/2 signalling pathway. A. Network depicting targets of Zfp263 across time. Shape of node represents type of target. Diamond shape represents TF, while circle represents gene. Edge color represents time point at which the Zfp263 binds to target (shown in Figure 4B). B. Venn showing number of targets in each time point and those shared across different combinations of time points. C. Table showing number of targets (genes/TFs) of Zfp263 across time. D. RNA-Seq gene expression values (scaled by row) of TFs from Figure 4A. E. Pathway analysis of Zfp263 targets (from Figure 4A) across time. F. Levels of phosphorylated ERK 1/2 were checked by western blot in WT and Zfp263 KO HoxB8 progenitor cells at day 4 of differentiation. Histone H3 was used as a loading control for the experiment. G. Levels of ERK and p-ERK (normalized to H3) were estimated from the western blot (A) by densitometry and plotted. Significance was tested by unpaired Welch’s T test (* p value ≤ 0.05, ** p value ≤ 0.01).

Chip-seq analysis on the HoxB8 neutrophils at days 1, 3, and 5 (**Supplementary Figure S5A**) found that Zfp263 was recruited within the 2500 base pairs to the TSS region of 148 out of the 192 direct “target genes”, indicating a strong concordance between the predicted and true binding sites for “target genes”. These were denoted by contouring the corresponding nodes of the network in black (**Figure 5A**). Zfp263 was recruited to all but one (Twist) of the identified “target TF genes”, either at a specific stage in neutrophil development (e.g. day 3 for Zfp263; day 3, day 5 for Tal1) or across all time points (Atp6v0c) **(Supplementary Figure S5B**). An interesting example of Zfp263 “target gene” included Zfp263 itself, implicating a feedback loop regulation, associated with fate-determining TFs (38). Tal2, Zbtb10, Maf, Maff are other TFs previously shown to impact developmental processes by influencing differentiation and proliferation of immune and non-immune cells in various tissues (39–41). In contrast, IRF7 and ETS2 are commonly linked to the development of inflammation (42–44). These TFs may contribute to regulation of distinct neutrophil transcriptional programmes downstream of Zfp263, but more work is needed to dissect their interactions.

Here we investigated the functional link between Zfp263 and the ERK1/2 signalling cascade, which is known to support neutrophil proliferation and block neutrophil maturation downstream of the G-CSF receptor (17, 19). The ERK1/2 signalling cascade was invariably highlighted by pathway analyses in this study (**Figure 1, S2, S4**), including of the Zfp263 “target gene” network (**Figure 5E**). We examined the protein expression levels of ERK1/2 and phosphorylated ERK1/2 (p-ERK1/2) in WT, Zfp263KO and Zfp263AP HoxB8 cells by western blot assay. The expression of ERK1 and p-ERK1 was substantially higher in the Zfp263KO cells post day 3 (**Supplementary Figure S5C, D**). More detailed inspection of ERK1/2 and p-ERK1/2 levels at day 4 of HoxB8 differentiation clearly demonstrated the reverse correlation between Zfp263 and ERK1/2 protein levels: depletion of Zfp263 resulted in higher levels of ERK1/2 and p-ERK1/2, whereas higher levels of Zfp263 led to inhibition of ERK1 levels (**Figure 5F, G**).

We thus concluded that Zfp263 and the ERK1/2 signalling pathway are functionally linked and hypothesised that Zfp263 may control neutrophil maturation via regulation of the ERK1/2 levels and activity.

### Zfp263 promotes neutrophil maturation by inhibiting the ERK1/2 signalling pathways

To confirm the causal Zfp263-ERK1/2-neutrophil maturation link, we tested if the inhibition of the ERK pathway. We treated HoxB8 progenitor cells with an ERK inhibitor or inhibitors of the upstream MEK kinases (P98059 and U0126) in increasing concentrations to confirm their effect on ERK phosphorylation. We observed that all the inhibitors were capable of reducing the phosphorylation of ERK1/2 on day 1 of differentiation but the dose response differed between WT and Zfp263KO differentiated cells (**Supplementary Figure S6A**). We then differentiated WT and Zfp263KO progenitor in the presence of either 50μM U0126 or 100μM ERK inhibitor, or 100μM P98059 and examined neutrophil maturation by flow-cytometry using the surface markers Ly6G and CD 101 across the time points-day 1, 3 and 5. We found that the ERK and MEK inhibitors were not effective in promoting the maturation of WT neutrophils, but were able to partially restore the delay in maturation of the Zfp263KO cells, increasing the percentage of mature neutrophils by ∼2 fold or more compared to the control (DMSO) condition (**Figure 6A, B**).

**Figure 6:**
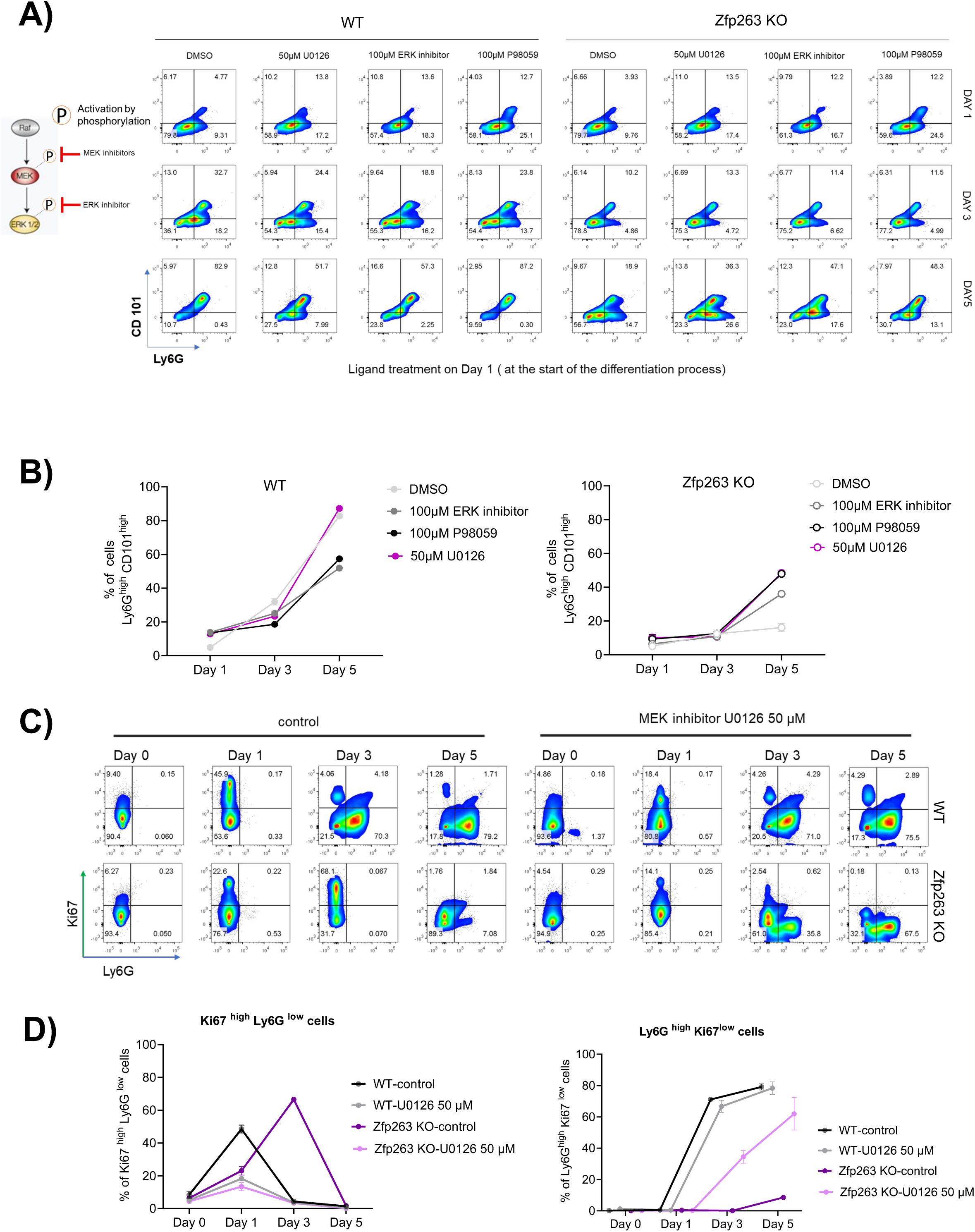
Treatment of Zfp263KO progenitors with ERK inhibitors restores neutrophil maturation. A. Differentiation of HoxB8 cells (WT and Zfp263 KO) was checked on Days 1,3 and 5 in the presence of 100 µM ERK inhibitor, 50 µM MEK inhibitor U0126 and 100 µM MEK inhibitor P98059. The inhibitors were added to the differentiation media on Day1 and expression of Neutrophil maturation cell surface markers Ly6G and CD101 were estimated by flow-cytometry. Ly6G^high^ CD101^high^ cells were gated to determine the % of mature neutrophils. B. The mean % of Ly6G ^high^ CD101^high^ cells from three technical replicates from Figure 6A have been plotted for WT and Zfp263 KO HoxB8 cells. C. Expression of Neutrophil differentiation cell surface marker Ly6G and proliferation marker Ki67 were estimated by flow-cytometry in HoxB8 cells (WT and Zfp263 KO) in the presence of MEK inhibitor U0126 (50 µM). Cells were differentiated and collected on Days 0, 1, 3 and 5 for the control (DMSO treated) and inhibitor treated conditions. D. The mean % of Ki67^high^ Ly6G ^low^ (left) and Ly6G ^high^ Ki67^low^ (right) cells have been plotted for Days 1,3 and 5 (from three technical replicates) as shown in Figure 6C.

Having observed that the levels of phosphorylated ERK1/2 showed a sharp increase in Zfp263KO cells post day 3 (**Supplementary Figure S5C, D**), we asked if the treatment of the cells with the ERK/MEK inhibitors on day 3 could be sufficient to restore the differentiation of Zfp263KO cells. We found that U0126 effectively increased the percentage of mature neutrophils among the Zfp263KO cells measured at day 5 (**Supplementary Figure S6B, C**). This indicates that the increase in ERK phosphorylation in Zfp263KO cells post-day 3 (later stages of differentiation) is linked to the delayed maturation of these cells.

We then used the well-established proliferation marker Ki67 to check if ERK1/2 inhibition led to a difference in the proliferation potential of the differentiating neutrophils. As the neutrophils mature, they show a higher Ly6G expression and a lower Ki67 (proliferation marker) (**Figure 6C, D**). Post day 3, the HoxB8 WT cells showed a significantly reduced Ki67+ population of proliferating cells, while the Ly6G+Ki67-population increased. Zfp263KO cells displayed a higher proportion of Ki67+ cells beyond day 3 and expectedly had a lower Ly6G+Ki67-population. The use of the MEK inhibitor U0126 was effective in reducing the Ki67+ population in the Zfp263KO cells (**Figure 6C, D**), confirming that higher ERK1/2 phosphorylation levels (post day 3) in these cells support cell proliferation at the expense of cell maturation.

Collectively, our results demonstrate that Zfp263 functions as a transcriptional repressor of the ERK1/2 signalling cascade in neutrophils, which limits their proliferative capacities and induces the process of differentiation into mature cells.

## Discussion

The process of granulopoiesis has long been studied as an example of changing cell morphology. Recent discoveries have added the repertoire of distinct cell surface receptors and specific gene expression profiles allowing for a better molecular definition of different developmental stages (13). These were further expanded by single-cell studies (9, 22, 27, 45, 46), which also demonstrated a high molecular similarity of neutrophil maturation stages across human, mice, and zebrafish species (47, 48). However, TF networks that promote the transition between stages of neutrophil maturation remain largely unknown. Here using the model system of controlled neutrophil maturation, we identified a network of putative global regulators acting across all stages of neutrophil development, with Zfp263 at its apex. We demonstrate that Zfp263 acts as a checkpoint of neutrophil maturation.

Zfp263 belongs to the Kruppel-like C2H2-type zinc-finger protein family, is conserved across species (in human, chimpanzee, and mouse) and has both the SCAN and KRAB domain (49). Despite the KRAB domain often being linked with transcriptional repression, a comprehensive analysis of Zfp263 targets in K562 and Hela cells revealed that Zfp263 may have a multifaceted role in regulating gene expression, capable of both positively and negatively influencing transcription (50). Interestingly, our analysis of gene expression in neutrophils deficient in Zfp263, indicated that it may act as an overall transcriptional repressor at the earlier stages and as transcriptional activator at the later stages of neutrophil maturation trajectory (**Supplementary Figure S7**), implying the possibility of Zfp263 interactions with distinct co-factors at different stages of neutrophil development, possible via its SCAN domain (50, 51). Consistently, we observed that gene categories associated with neutrophil effector functions and later stages of maturation, such as cytokine and chemokine production, were under positive regulation by Zfp263 (**Supplementary Figure S4**).

The relationship between Zfp263 and ERK1/2 signalling presents an intriguing mechanism by which Zfp263 may regulate neutrophil maturation. The observed increase in ERK1/2 and p-ERK1/2 levels in the Zfp263 KO cells at the later stages of neutrophil differentiation suggests that Zfp263 acts as a transcriptional repressor of the ERK1/2-driven proliferation and facilitator of neutrophil differentiation. This hypothesis is supported by the partial restoration of neutrophil maturation in the Zfp263 KO cells treated with ERK/MEK inhibitors (**Figure 6**). In addition, earlier studies demonstrated that prolonged activation of ERK1/2 leads to augmented activation of c-Fos and Egr1, which is needed to tilt the balance in the myeloid precursor development towards monocytes, while inhibition of ERK1/2 favours neutrophil development (19). Thus, it is exciting to speculate that in addition to its function as a checkpoint of neutrophil maturation, Zfp263 may also act as a gatekeeper of neutrophil-monocyte balance.

The Zfp263 “target gene” network contained several previously described transcriptional regulators of neutrophil maturation, including Klf6 and Gfi-1 (23, 26). Gfi-1, a zinc finger TF, ensures the differentiation of GMPs into neutrophil granulocytes by blocking enhancers and expression of genes that promote myeloid differentiation (52). While expression of Gfi1 decreases during neutrophil differentiation and is absent in mature neutrophils (27, 52), expression of Zfp263 has the opposite trend (**Figure 2**). Zfp263 deficient cells display higher levels of Gfi1 (**Figure 5**), thus, it is likely that Zfp263 acts as a transcriptional repressor of Gfi1, but more work is needed to dissect their interactions. Of interest, G-CSF signalling was found to be impaired in the absence of Gfi1, due in part to a deregulation of RASGRP1 upstream of the ERK1/2 pathway (53). Therefore, it is possible that interactions of Gfi1 and Zfp263 may also exist within the ERK1/2 pathway regulation. Zfp263 deficiency also impacted core neutrophil immune and effector functions, such as (trans)migration, cytokine, ROS and NET production (**Figure 4**). Some of the direct targets of Zfp263, may in fact be critical drivers of immune functions. For example, IRF7 expression was abundant in neutrophils from patients with severe COVID-19 and correlate with neutrophil-mediated inflammatory responses (54). The temporal interactions between Zfp263 and target TFs remain to be elucidated, as well as the possibility of uncoupling the neutrophil developmental and functional programmes.

While the role of Zfp263 in neutrophil biology has never been explored, a number of recent cancer studies highlighted its possible role in tumorigenesis (55–60). In lung adenocarcinoma, its high expression was associated with a better clinical response to the EGFR inhibitors. This was partially due to its ability to directly suppress the EGFR expression (59). On the contrary, high Zfp263 expression in human pancreatic ductal adenocarcinoma, was linked to poor prognosis (60). Our own analysis of Zfp263 expression levels in cancer datasets from the Cancer Genome Atlas (TCGA), highlighted a number of cancers where Zfp263 showed a significant positive effect on survivability based on the Kaplan-Meier formula, including Acute Myeloid Leukemia (LAML) and Ocular melanomas (UVM), Adrenocortical Cancer (ACC), but also detected a negative correlation with Prostate Cancer (PRAD) (**Supplementary Figure S8**). Further exploration into Zfp263 function will need to be conducted in the context of specific cancer types and immune cell involvement.

In summary, our study reveals the key dynamics of chromatin accessibility and gene expression programmes that orchestrate neutrophil maturation, with Zfp263 emerging as the key transcription factor. We showed that Zfp263 is a transcriptional checkpoint of neutrophil maturation in the balance between proliferation and maturation. The insights gained from mapping out a new regulator of neutrophil maturation not only advances our understanding of neutrophil biology but also present new opportunities for therapy in diseases characterized by aberrant granulopoiesis, such as cancers and other inflammatory disorders. Future research should explore the therapeutic potential of targeting Zfp263 and its downstream pathways, aiming to modulate neutrophil development and function in disease. In fact, its overexpression via adeno-associated viruses has been shown to suppress tumour growth in lung adenocarcinoma models (59). Additionally, elucidating the interactions between Zfp263 and other transcriptional regulators within the neutrophil maturation network will be crucial for comprehensively understanding the control mechanisms at play and how they might be leveraged for therapeutic benefit.

## Methods

### Generation of HoxB8 neutrophils with Zfp263 knockout

The HoxB8 neutrophils generation was done like our previous protocol (23, 61). Briefly, to generate CRISPR-Cas9-mediated Zfp263 knockout, progenitors were transduced with TLCV2 lentiviruses targeting exon 1 of Zfp263 (ENSMUSG00000022529; guide RNA1 (gRNA1): CACCGGCTCCTCATAGTGAAACTGG; gRNA2: AAACCAGTTTCACTATGAGGAGCC). Briefly, lentivirus containing our gRNA of interest was produced from HEK-293T cells transfected with the gRNA-containing plasmid mixed with the packaging vectors pVSVG (plasmid no. 8454; Addgene) and psPAX2 (plasmid no. 12260; Addgene). The lentivirus-containing supernatants were collected 48h post-transfection, and subsequently, every 24h for three days. For lentivirus transduction, HoxB8 cells were cultured with virus containing-media and polybrene for cell membrane permeability. 72h after transduction, the cells were cultured with 6 µg/mL puromycin for antibiotic selection of the targeted Zfp263 knockout.

#### Culture and differentiation of HoxB8 cells

HoxB8 progenitors are maintained in complete RPMI-1640 medium supplemented with 30 μM of β-mercaptoethanol, 4% SCF-containing supernatant and 10 uM estradiol in a 5% CO2 tissue culture incubator at 37 °C. The differentiation of the HoxB8 myeloid progenitors was induced by estradiol removal and culture in complete RPMI-1640 medium supplemented with 30 μM of β-mercaptoethanol, 4% SCF-containing supernatant and 20 ng/mL G-CSF in a 5% CO2 tissue culture incubator at 37 °C.

### RNA-seq analysis

We used FastQC to assess the quality of reads from the bulk RNA-seq experiments (62). Trimmomatic was then used to remove adapter and low-quality sequences (63). We then used STAR to align the reads to the mouse reference genome (mm10) and generate aligned & sorted BAM files (64). HT-seq was used to count the mapped reads from the BAM files (65). The count matrix generated from HT-seq was normalized in R using the DESeq2 package (66). Differential expression analysis was performed using Wald’s test in the DESeq2 package. The weighted gene correlation network analysis (WGCNA) was used to identify gene modules that correlate with maturation trajectory and knock-out (67). WGCNA was also be used for the construction of module-specific transcription factor (TF) to target correlation networks. Network properties such as module membership and degree were used to filter the TF-Target network(67). Gene modules was annotated using gene ontology analysis and Reactome pathway analysis available in the clusterProfiler package in R (32).

### ATAC-seq, MOTIF analysis, and Zfp263 regulatory network

Raw reads were trimmed using Cutadapt v2.9 (68) and then aligned to the mm10 mouse reference genome using Bowtie2 (69). Picard tools v2.22.3 was used to remove PCR duplicates. Before peak calling all the reads that mapped to chrM, with a mapping quality <10 and insert sizes >150bp were removed. Model-based analysis of ChIP-seq (MACS2) program was used for peak calling and merging (70). Merged peaks was counted using BedTools (71). The read counts were input to R, DESeq2 was then used for identifying significant DAPs (p_adj_ < 0.05) using likelihood ratio test. For time-based comparisons, significant DAPs (p_adj_ < 0.05 and fold change > 1) were identified using Walds.

The MEME-Chip suite was used for performing de novo motif discovery on the differentially accessible peaks (36). The motif database from JASPER, HOMER, and HOCOMOCO was used to identify enriched motifs and perform transcription factor binding site analysis (72–74). Based on the presence of TF-motif from the motif analysis, a TF-Target network was constructed for all DAPs.

The Zfp263 network was constructed by identifying genes with DAPs within the 2,500 base pair regions proximal to the transcription start site that also contained the Zfp263 binding motif, based on the de novo motif discovery using the MEME-Chip suite. Further filtering retained only those genes that were differentially expressed (in the RNA-Seq data) between WT and Zfp263KO at the respective time point. Consequently, the network was annotated for specific time points, depending on the presence of the DAP, the Zfp263 motif, and DEG at those respective times.

### Chromatin immunoprecipitation (ChIP)

20 million HoxB8 (WT and Zfp263 KO) cells at various stages of neutrophil differentiation (Day 0, 1, 3, and 5) in 50 ml of cell culture media were fixed for 10 mins at RT (room temperature) using 1% Formaldehyde (final concentration) and the reaction was quenched using 125 mM Glycine. The cells were washed thrice with chilled 1X PBS, then incubated for 10 mins in lysis buffer 1 on ice and pelleted at 1350 x g for 5 minutes at 4°C. The supernatant (cytoplasmic fraction) was removed and the pelleted cell nuclei were resuspended in lysis buffer 2 and incubated at RT for 10 mins. Cell nuclei were pelleted at 1350 x g for 10 minutes at 4°C resuspended in lysis buffer 3 and incubated for 5 mins on ice. The cell nuclei were sonicated for 10 cycles (30 seconds -ON and 45 seconds OFF) on the Diagenode Bioruptor Pico instrument while maintaining the water bath temperature at 4°C.

5% of the sonicated faction (1ml) was used to isolate DNA using Qiagen PCR purification kit and checked for an optimal post-sonication mean fragment size of 200-250 bp using 1.5% Agarose gel (in 1xTAE) electrophoresis. The fragment size was further checked using the Agilent tape-station capillary electrophoresis platform (D1000 kit) in standardisation experiments.

10% of the sonicated fraction was stored at -20°C to be used as the input fraction. The remaining fraction was used to aliquot two 400 µl fractions. In one fraction, 10 µg of Zfp263 antibody (100 µl of 0.1 µg/µl primary antibody) was added. In the other fraction - 10 µg of IgG(mock) antibody was used and volume was made up to 500 µl using lysis buffer 3. 100 µl Dynabeads-protein G (ThermoFisher) were added to both fractions followed by gentle mixing overnight on a rotator at 4°C. Following this, the tubes containing the fractions were placed on a magnetic rack and the supernatant was removed. Low salt buffer was added to pelleted beads on the magnetic rack and put on a rotator at 4°C for 5 mins. This was repeated two more times and then the same washing steps were repeated for high salt and LiCl buffers. The beads were resuspended in 100 µl of Elution Buffer at RT and incubated with 10 µl of proteinase K (Invitrogen™ AM2546-20 mg/ml) at 65°C for 2 hours. At this stage the input DNA was thawed and incubated at 65°C with proteinase K. The DNA was purified from all the samples using Qiagen PCR purification kit and resuspended in 50 µl of Nuclease-free water at RT.

In all the above steps, 1 ml of the buffers were used for resuspension and in steps following formaldehyde fixation and the buffers used were prechilled to 4°C unless otherwise mentioned. All the buffers used at 4°C had 2X mammalian protease inhibitor cocktail (Sigma-P8340) added to it. The Buffer compositions are enlisted in Supplementary Method Table 1.

### ChIP-sequencing

The samples were quantified by Qubit dsDNA HS kit using the manufacturer’s recommended protocol and absorbance (A) ratio (A 260nm/A 280nm) was estimated by Nanodrop one (ThermoFisher) Spectrophotometer. It was ensured that A 260/280 ratios for samples were within 1.8-2 and a minimum of 14 ng of DNA (per sample) was sent for preparation of the ChIP-seq libraries at the Oxford Genomics Centre (University of Oxford). The TruSeq ChIP Library Preparation Kit (Illumina) was used to make libraries that were sequenced on the NovaSeq 6000 platform from Illumina. The raw FASTQ files were used for sequencing QC and downstream analysis.

### ChIP-seq analysis and validation of Zfp263 regulatory network

FastQC was used for performing quality checks on the raw data (62). Trimmomatic was used to remove the adapter sequences from the reads (63). Bowtie2 was then used to align the reads to the mouse genome (mm10)(69). Samtools was used to convert, and filter aligned reads to BAM (75). MACS2 program was then used for peak calling (70). The peaks of the replicates were then combined using IDR and BEDtools (71). Both de novo motif discovery and known motif enrichment was then done using the MEME-suite (36). The results from the ChIP-seq were used to validate the Zfp263 regulatory network, Figure 5A.

### ChIP-qPCR

Enrichment of Zfp263 at specific sites in the genome was probed using custom-designed primer pairs using Sybr green dye-based detection of amplification products on the Quantstudio 7 (Thermo Fisher). The reaction condition and primer details are in the Supplementary Method Table 2. Enrichment was estimated over 1% input (2^ - (ct _sample_ - ct _input_)) for individual ChIP samples (Zfp263 IP and Mock IgG respectively). Fold change of Zfp263 enrichment over corresponding IgG (mock) enrichment was plotted for each sample.

### Western Blotting

1 million cells were washed with chilled 1X PBS thrice and resuspended in 200 µl 1X RIPA Buffer (with complete protease and Phospho-STOP phosphatase inhibitor cocktails from Sigma in recommended concentrations). The cells were kept on a rotator for 2 hours at 4°C to allow for lysis.

The cells were centrifuged at 13000 rpm for 10 minutes at 4°C and the supernatant was collected. The protein was quantified using the Pierce BCA protein assay (ThermoFisher) kit as per the manufacturer’s recommended protocol. 20-40 µg of protein was denatured with a final concentration of 1X LDS sample buffer (Life Technologies) at 95°C for 10 mins. Following this, the protein was resolved on NuPAGE-4 to 12%, Bis-Tris Gels (Life Technologies) in 1X MOPS/MES Buffer (NuPAGE-Life Technologies). A broad-range (12-225 Kda) protein marker (Amersham ECL Rainbow -RPN800E) was run along with the samples for molecular weight reference. The protein was transferred onto a Nitrocellulose membrane using the Trans-Blot Turbo semi-dry transfer setup as per the manufacturer’s recommendations. The transfer was checked with Ponceau staining solution (Life Technologies) and destained with 1X PBS, followed by blocking the membrane with 5% BSA (in 1XPBS) at RT for 2 hours. The membrane was then washed with 1X PBS (for 10 minutes) thrice and incubated with primary antibody (0.5-1 µg /ml in 1% BSA) for 2 hours at RT or overnight at 4°C. Subsequently, the membrane was washed with 1X PBST (1X PBS with 0.1% Tween-20) thrice for 10 minutes and incubated with appropriate HRP (horse-radish peroxidase) tagged secondary antibody (1:2000 dilution (in 1% BSA) for 2 hours at RT. While probing for phospho-proteins, PBS was replaced with TBS in all the buffers used. The membrane was washed thrice with 1X PBST for 10 minutes each and developed using Clarity-max chemiluminescent HRP substrate (BIO-RAD) using the manufacturer’s protocol. The chemiluminescent signal from the membrane was captured using the Syngene G-box gel imaging instrument.

### Flow-cytometry

To prepare cells for flow cytometry, 1-2 million cells were washed in 1X chilled PBS and stained with live/dead staining dye (L34976-Life Technologies) as per the manufacturer’s staining recommendations. Following this, the cells were washed thrice with chilled FACS Buffer (PBS + 0.1% BSA + 0.01% sodium azide + 1mM EDTA). The cells were incubated using Fc Block (BD Biosciences) (1:100 in FACS buffer) for 10 mins at 4°C and then stained with fluorophore-conjugated cell surface marker antibodies (1:200) for 1 hour at 4°C in the dark. The cells were washed with FACS buffer thrice and fixed with CytofIx buffer (BD).

For intercellular staining, cells were permeabilised with Foxp3 Fix/Perm Buffer (eBioscience™ Foxp3 / Transcription Factor Staining Buffer Set 00-5523-00) and then were similarly blocked and stained with the intra-cellular antibody as stated above. The Flow cytometry data was acquired using the FACSDiva v.6.1.3 software on the LSRFortessa X-20 flow cytometer (BD Biosciences). Data analysis was done using FlowJo v.10 (FlowJo LLC).

### Antibodies

The antibodies used have been listed in Supplementary Method Table 3.

### ERK inhibitor treatment

Commercially available ERK (MercK-328006-5MG) and MEK inhibitors from Abcam (PD 98059 (DMSO solution), MEK1 inhibitor; U0126, Selective MKK inhibitor) were procured as used for treating cells as indicated in individual Figure legends.

### In vivo experiments

Prior to in vivo adoptive transfer, HoxB8 progenitors were washed twice in HBSS and live cells were counted using hemacytometer and Erythrosine B staining. B6.SJL-Ptprc^a^Pepc^b^/BoyJ recipient mice expressing the CD45.1 leukocyte surface marker aged between 69 and 152 days from both sexes were irradiated using a ^137^Cs source with a Gamma-Service Medical D1 irradiator at 11 Gray for 17 minutes and injected retro-orbitally the same day with 5 × 10^7^ HoxB8 progenitors in 200 μl HBSS. Mice were kept in individually ventilated cages in Semmelweis University Basic Medical Science Center’s minimal disease animal facility on oral antibiosis (300 mg/ml neomycin, 6 mg/ml polymyxin B). 5 days after progenitor injection, 10 μl blood was drawn from recipients followed by the i.p. injection of 1 ml sterile thioglycolate (Sigma STBMFTM12). 4 hours later, blood was sampled again and mice were sacrificed by cervical dislocation under isoflurane anesthesia. Peritoneal infiltrate was collected by washing the abdominal cavity with 5 ml ice cold flow buffer (PBS, 1% BSA, 5 mM EDTA and 2 v/v% heparin, pH = 7.4), filtered through 70 μm cell strainer and washed twice in PBS. BM was isolated from 2 femora per animal by flushing the bones with 5 ml flow buffer, clumps were disaggregated by thorough pipetting followed by filtration through 70 μm strainer. All samples were incubated in Mouse BD Fc Block^TM^ (5 μg/ml in flow buffer, BD 553142) for 5 minutes on ice followed by staining for flow cytometry using the following antibodies for 40 minutes on ice in flow buffer: Ly6C FITC (1.25 μg/ml, BD 553104), B220 PE (0.5 μg/ml BD 553089), Siglec F PE (0.5 μg/ml, BD 552126), F4/80 PE (0.5 μg/ml, BD 565410), CD45.1 biotin (1.25 μg/ml, BD 553774), CD101 PE-Cy7 (0.3 μg/ml, Invitrogen 25-1011-82), Ly6G PerCP-Cy5.5 (0.5 μg/ml, BioLegend 127616), CD11b AlexaFluor 647 (0.5 μg/ml, BD 557686). Samples were then washed and stained for 40 minutes on ice in PBS using 1:400 diluted LIVE/DEAD™ Fixable Violet Dead Cell Stain (Invitrogen L34964) and PE streptavidin (1.25 μg/ml, BD 554061). Red blood cells were lysed using a 150 mM NH_4_Cl lysis buffer and samples were measured immediately after staining without fixation on a CytoFLEX cytometer (Beckman Coulter). Among live singlet events HoxB8-derived neutrophils were identified as CD45.1^neg^ B220^neg^ Siglec F^neg^ F4/80^neg^ CD11b^pos^ Ly6G^pos^ events. Experiments were approved by the Animal Experimentation Review Board of Semmelweis University.

### Morphological assessment by cytospin

Morphological assessment was performed as described in our previous work (61). Briefly, differentiated neutrophils are pre-washed with dPBS, and spun onto glass slides by centrifugal forces using a Cytospin Cytocentrifuge (Thermo Fisher Scientific) at 400 g for 5 min. The slides were stained using the modified Wright–Giemsa stain Epredia™ Shandon™ Kwik-Diff™ Stains (Fisher Scientific) according to the manufacturer’s instructions and imaged using the brightfield Olympus BX51 fluorescence microscope (Olympus).

### Measurement of cytokine and chemokine production ELISA

The levels of TNF and CCL2 were quantified using the Mouse DuoSet ELISA (#DY479, R & D systems) following manufacturer’s instructions. The optical density of each well was quantified at 450nm wavelength on a SPECTROstar nano spectrophotometric Plate Reader (BMG Labtech) and corrected for plate absorption at 540nm. The data was exported by MARS Data Analysis software (BMG Labtech) and analysed in GraphPad.

### NET formation

To induce NETosis, 100 uL HoxB8 neutrophils (1 × 10^6^ cells/mL) were seeded into an 8-well Nunc Lab-Tek II Chamber Slide (VWR international) coated with 2% poly-L-lysine (MilliporeSigma). Neutrophils were stimulated with 10 μM of ionomycin and PMA (Merck, Sigma-Aldrich) or dPBS as control overnight at 37 °C in a 5% CO_2_ tissue culture incubator. Samples were subsequently fixed with 4% paraformaldehyde (Sigma-Aldrich) in dPBS for 30 min at RT. After blocking, the primary antibodies rabbit anti-citrullinated histone 3 (catalogue no. ab5103; Abcam) and mouse anti-mouse MPO (catalogue no. HM1051BT; Hycult) were added at a 5 ug/mL and 0.5 ug/mL respectively, and incubated overnight at 4 °C. Cells were washed with dPBS before adding secondary antibodies: mouse anti-rabbit DyLight 647-conjugated secondary antibody (Thermo Fisher Scientific) and rabbit anti-mouse IgG secondary antibody conjugated with Alexa Fluor 488 (Thermo Fisher Scientific). Antifade gold DAPI reconstituted mounting medium (Invitrogen) was used for nuclear staining and sealing. Images were obtained using a confocal ZEISS 980 fluorescence microscope. Neutrophils with a clear formation of fibers stained by citrullinated histone 3, colocalized with a diffuse nucleus stained by DAPI and colocalization with MPO, were counted as neutrophils under NETosis. Images were analysed using Fiji.

### Extracellular ROS production

As previously described in our previous work (14), extracellular ROS measurement was performed by a fluorometric assay–based method using OxyBURST H2 HFF Green BSA (Thermo Fisher Scientific). 100 uL HoxB8 neutrophils resuspended at a concentration of 1 x 10^7^ cells/mL KRP buffer (Merk) supplemented with 1.0 mM Ca^2+^, 1.5 mM Mg^2+^, and 5.5 mM glucose (pH 7.4). 10 μL of OxyBURST H2 HFF Green BSA was added to each well for 2 minutes at 37°C and treated with 1 μM fMLP. The release of extracellular ROS was measured for over 120 min, with 5 min intervals, with the FLUOstar Omega microplate reader (BMG Labtech). Sample was excited at 488 nm and detected at 530 nm.

### Data availability

The RNA-seq, ATAC-seq, and ChIP-seq data of the HoxB8 neutrophils at different timepoints are available upon request.

## Acknowledgement

This work was in part funded by the Hungarian National Research, Development and Innovation Office (KKP-129954 to Attila Mócsai) and the HUN-REN Hungarian Research Network (0207007 to Attila Mócsai).

## Supplementary figure legends

**Supplementary Figure S1:**
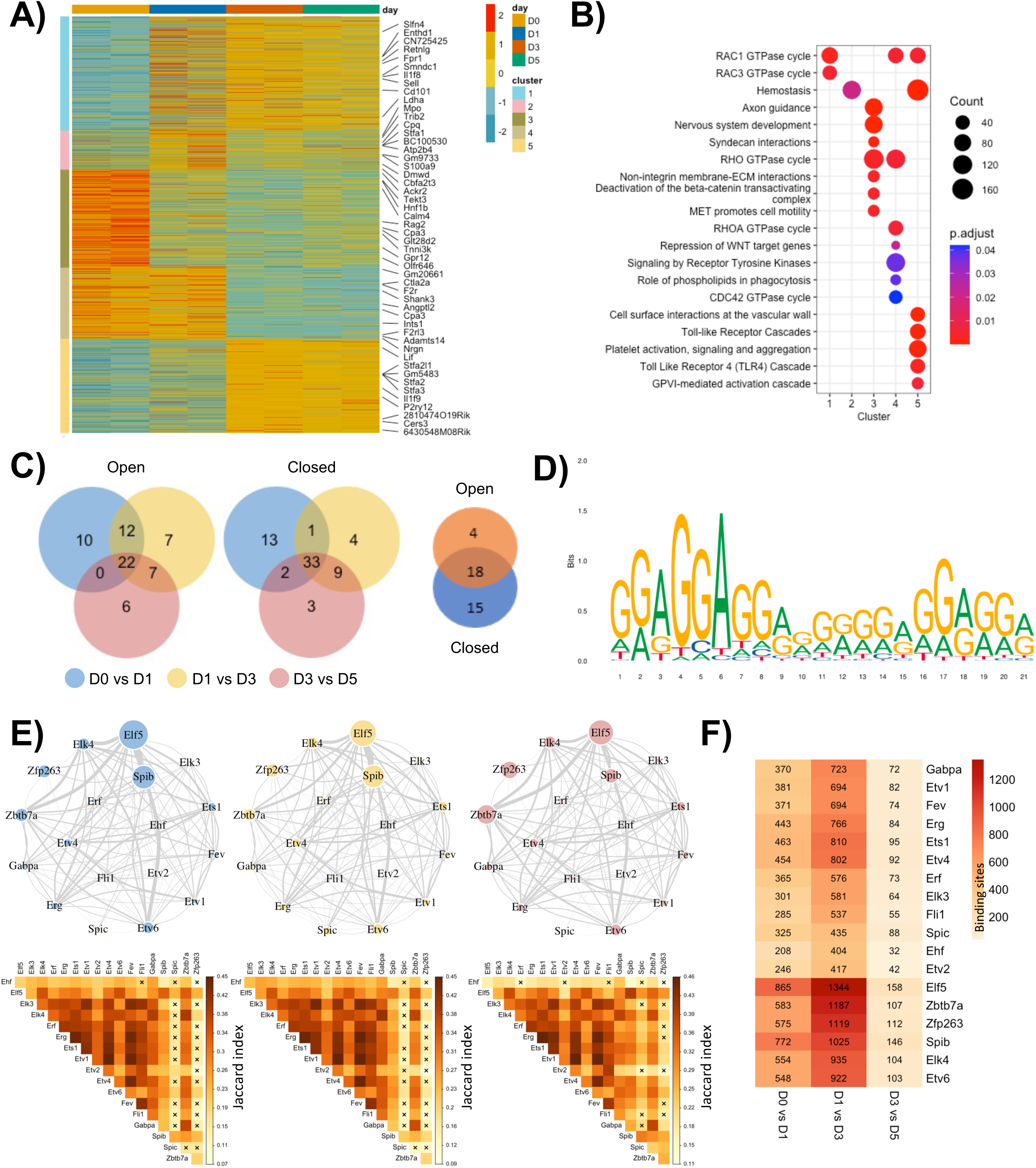
Changes in neutrophil chromatin landscape during their maturation and associated pathways and TFs. A. Heatmap representing distinct day-specific chromatin accessibility clusters. Clusters were obtained by hierarchical clustering of all DAPs (P_adj_ <0.05), based on Manhattan distances using Ward method. Each row in the heatmap is normalized to its minimum and maximum read counts in fragments in FPKM (fragments per kilobase of transcript per million mapped reads). B. Pathway enrichment analysis of the clusters from Supplementary Figure S1A. All pathways shown had a P_adj_ <0.05. C. Venn diagram showing enriched TF motifs across different comparisons between time points for open (left), closed (middle), and overlapping between open and closed (right). D. Zfp263 motif taken from Jasper database (motif id: MA0581). E. Top row represents the regulon analysis of the key transcription factors (as shown in Figure 1F) for each time point comparison shown in Supplementary Figure S1C. Size of the nodes represents number of binding sites detected based on the MEME suite analysis in the DAPs proximal to TSS, across all time points. Thickness of the edges represents the common number of binding sites shared between the corresponding TFs. Bottom row represents heatmap showing the Jaccard’s similarity index for the binding sites shared between different TFs for the corresponding time point. The **x** represents TF pairs that a significantly different (Fisher’s exact test; p-value <0.05) in terms of the shared binding sites. F. Heatmap quantifying the number of binding sites of each TF (in row) for the time point comparison (in column).

**Supplementary Figure S2:**
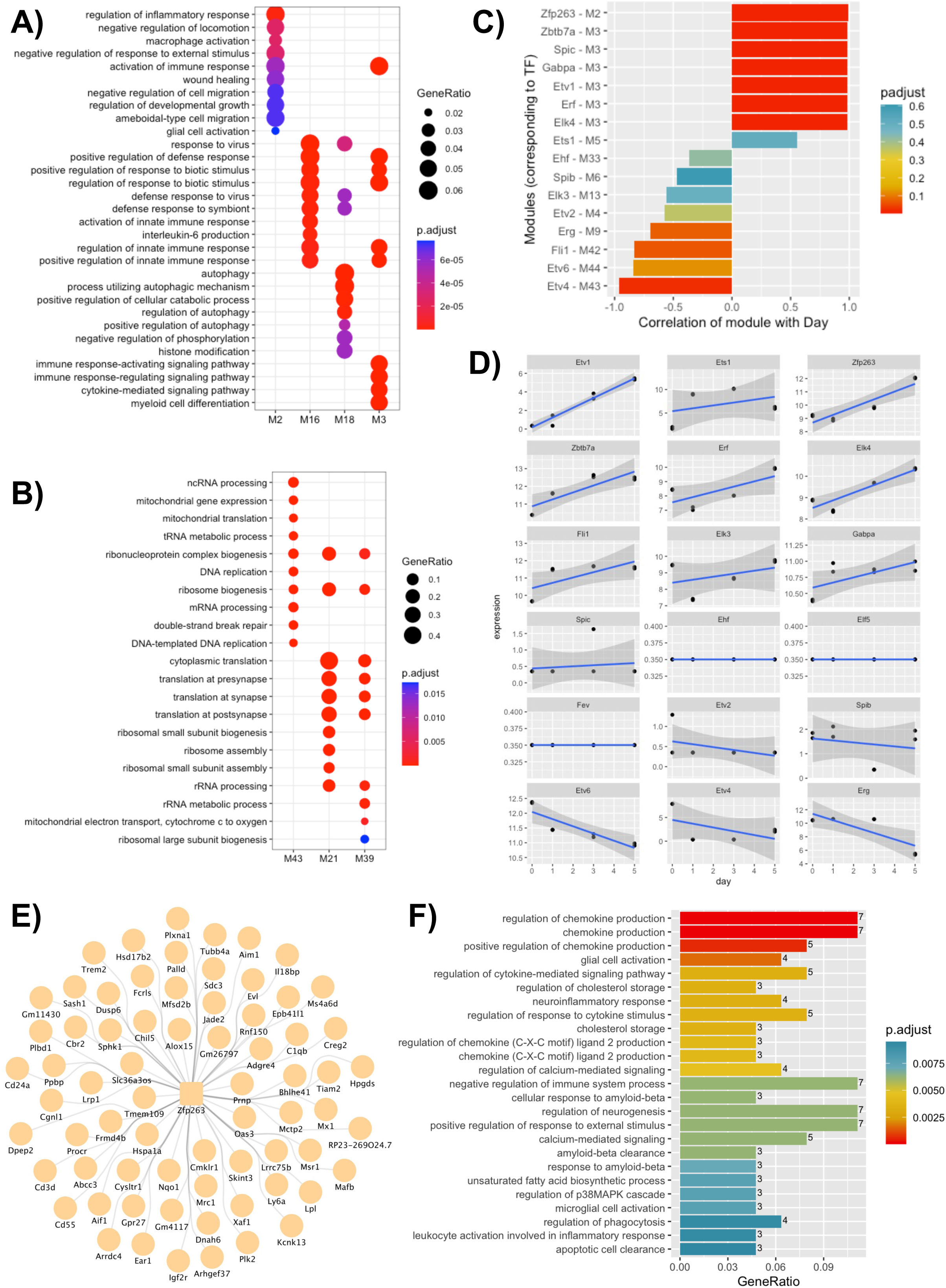
Gene expression profiling of HoxB8 neutrophils over days of maturation identifies modules and pathways with time-specific trends. A. Pathway analysis of modules that positively correlate (P_adj_ <0.05) with time. B. Pathway analysis of modules that negatively correlate (P_adj_ <0.05) with time. C. Pearson correlation between module mean eigen gene value vs. time. Here modules are chosen based on whether they contain the TFs in Figure 1G. D. Gene expression (y-axis) trend of all TF’s (in Figure 1G) with time (x-axis). E. Zfp263 correlation network. The network comprises of genes in the M2 module that highly correlate in gene expression with Zfp263. F. Pathway analysis of the Zfp263 correlation network (Figure S2E)

**Supplementary Figure S3:**
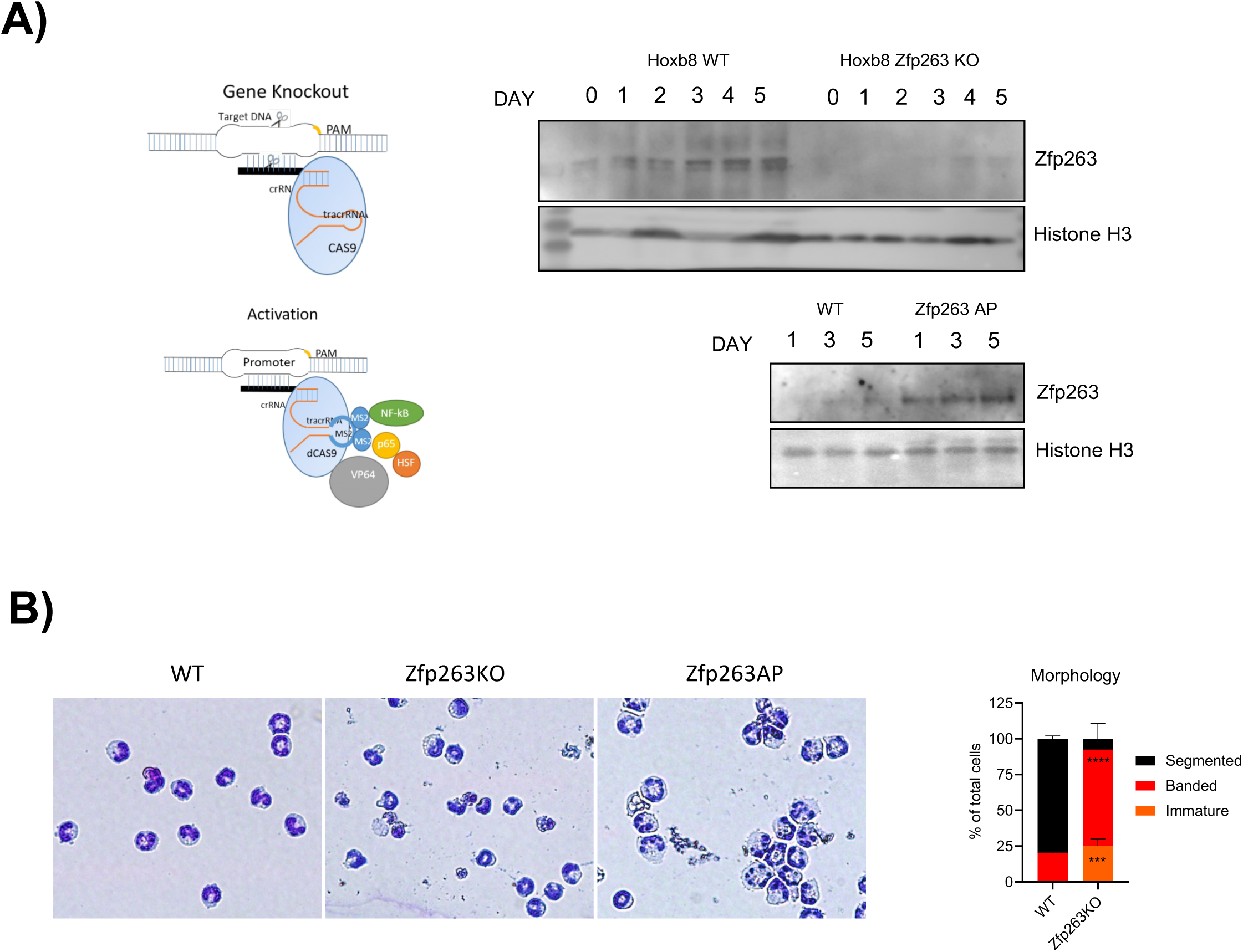
ZPF263 depletion delays neutrophil morphological maturation, while its activation expedites it. A. HoxB8 cells with ZPF263 depletion (Zfp263 KO) or activation (Zfp263 AP) were generated by CRASPR-Cas9 mediated gene editing as shown in the schematics (see methods for details). ZPF263 expression was checked by western blot to confirm depletion of the protein in Zfp263 KO cells (top) and enhanced expression in Zfp263 AP cells (below) at various time points of differentiation as indicated. Histone H3 was used as a loading control. B. Morphological assessment WT, Zfp263KO, and Zfp263AP neutrophils using Wright-Giemsa staining. For quantification - the results are expressed as percentages of segmented, band neutrophils and metamyelocyte of at least 200 cells counted from different fields. Statistical comparison was made by two-way ANOVA, ***P < 0.001 and ****P < 0.0001. Data are shown as means and SD from three independent experiments.

**Supplementary Figure S4:**
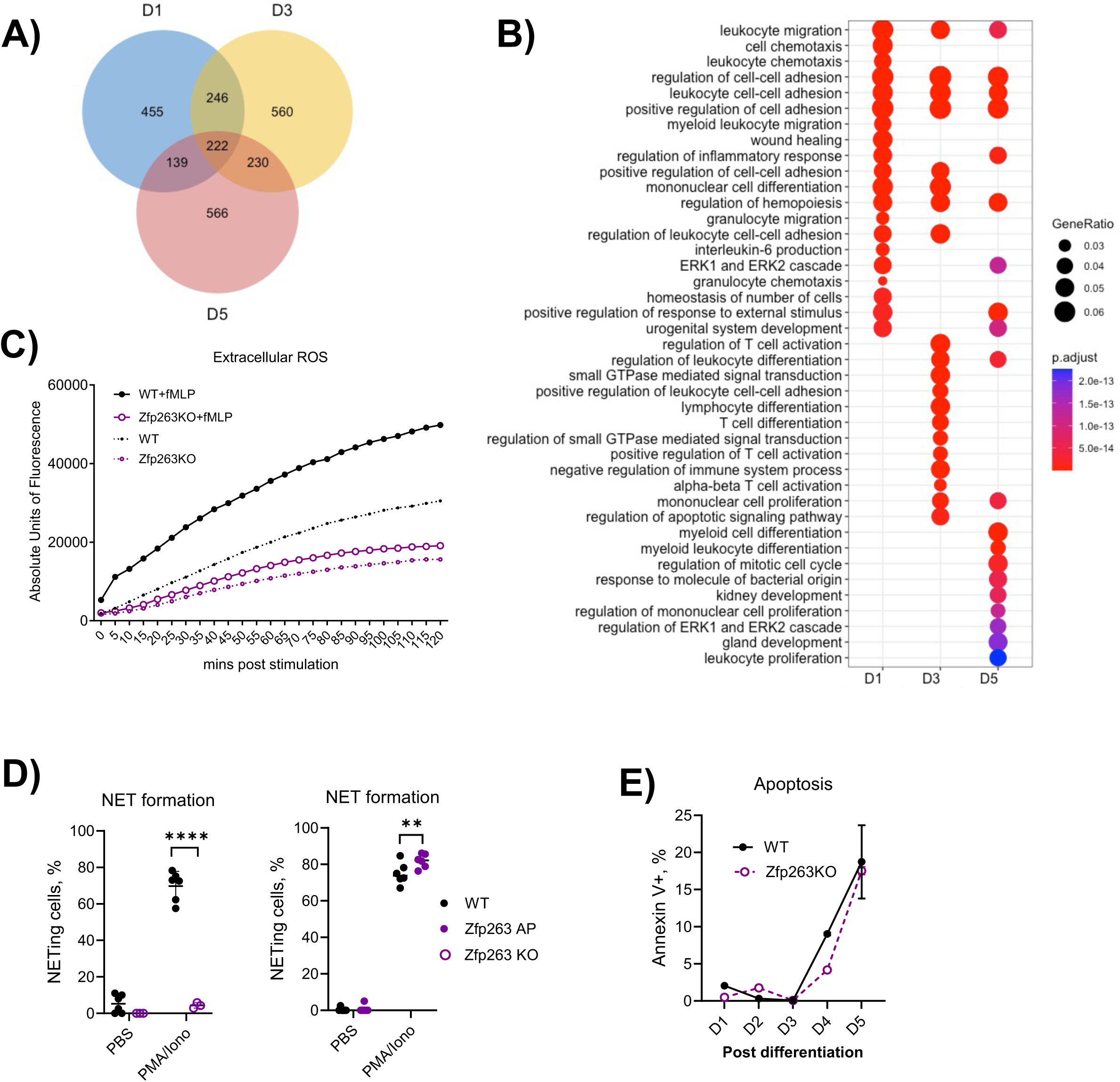
Combined analysis of differential gene expression and chromatin remodelling highlights Zfp263 dependent signalling and inflammatory pathways. A. Venn diagram showing number of genes that are positively correlated between ATAC and RNA-Seq data in Figure 4D (red dots) for day 1, 3, and 5. B. Pathway analysis of all genes as shown in Supplementary Figure S4A. C. Quantification of extracellular ROS production by WT and Zfp263KO HoxB8 neutrophils with and without fMLP-induction. Data are from a representative experiment. D. NET formation in response to stimulation by PMA/ionomycin in WT and Zfp263KO (left) or Zfp263AP HoxB8 neutrophils. Data are expressed as the percentages of neutrophils undergoing NETosis out of at least 200 cells counted from different fields and independent replicates. Statistical comparison was made by two-way ANOVA, **P<0.01, ****P<0.0001. Data are shown as means and SD from four independent experiments. E. Percentage of Annexin V+ cells at different days of differentiation in WT and Zfp263 KO Hoxb8 cells. Data are shown as means and SD from three independent experiments.

**Supplementary Figure S5:**
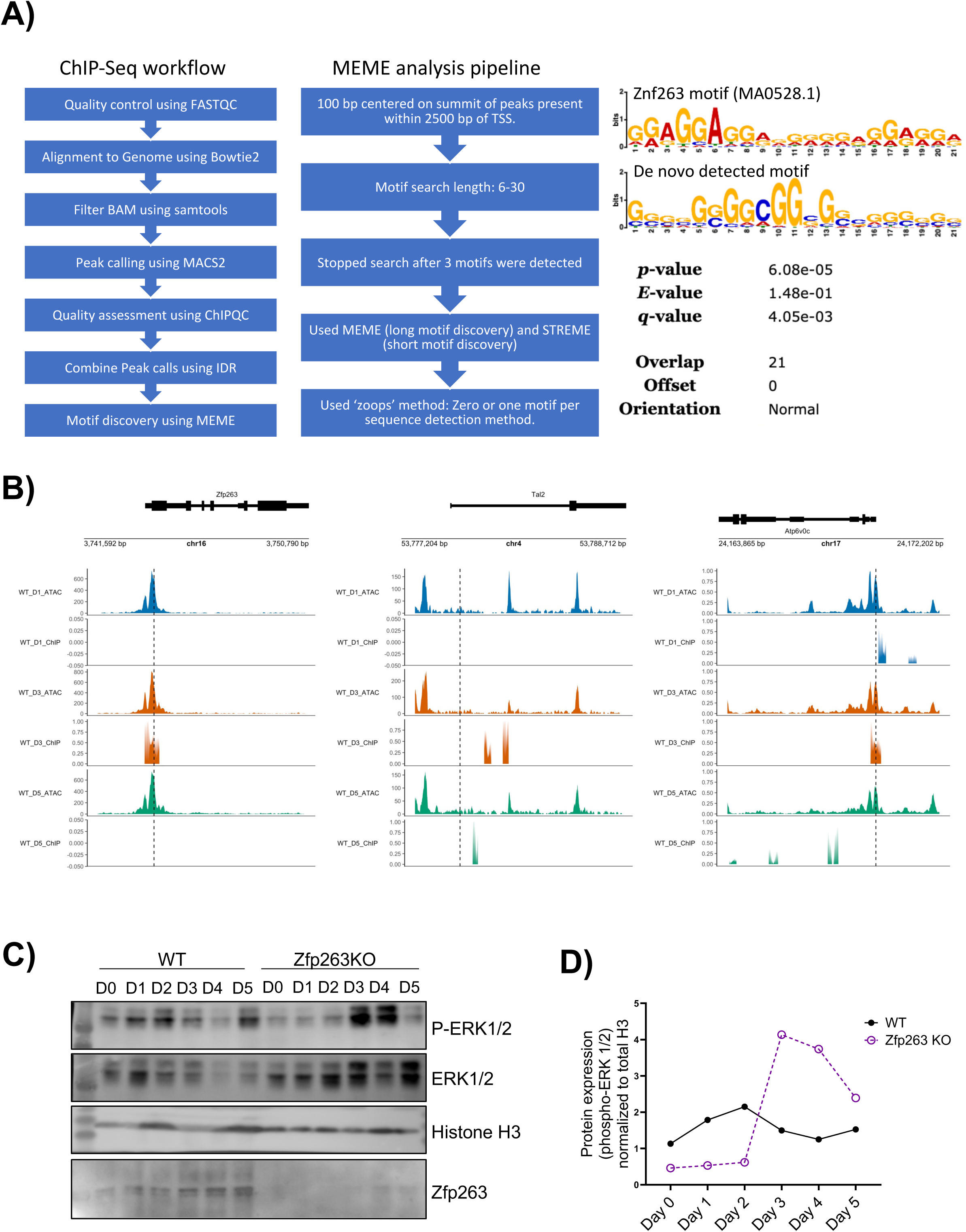
ChIP-seq analysis a strong concordance between the predicted and true Zfp263 binding sites. A. Steps involved in ChIP-Seq analysis pipeline (left panel), MEME analysis (center panel), and the de novo motif of Zfp263 discovered from ChIP-Seq data is significantly similar to the Zfp263 motif (MA0528.1) from Jasper database. B. ATAC-Seq and ChIP-Seq peaks of Zfp263, Tal2, and Atp6v0c. Dashed line represents transcription start site (TSS). C. Levels of Total ERK 1/2 and phosphorylated ERK 1/2 were checked by western blot in HoxB8 progenitor cells (WT and Zfp263 KO) for days 0,1,2,3,4and 5 of differentiation. D. Levels of phosphorylated ERK 1/2 were plotted for HoxB8-ER progenitor cells (WT and ZNF263 KO) by densitometry analysis of the western blot experiment as shown in Figure S5C.

**Supplementary Figure S6:**
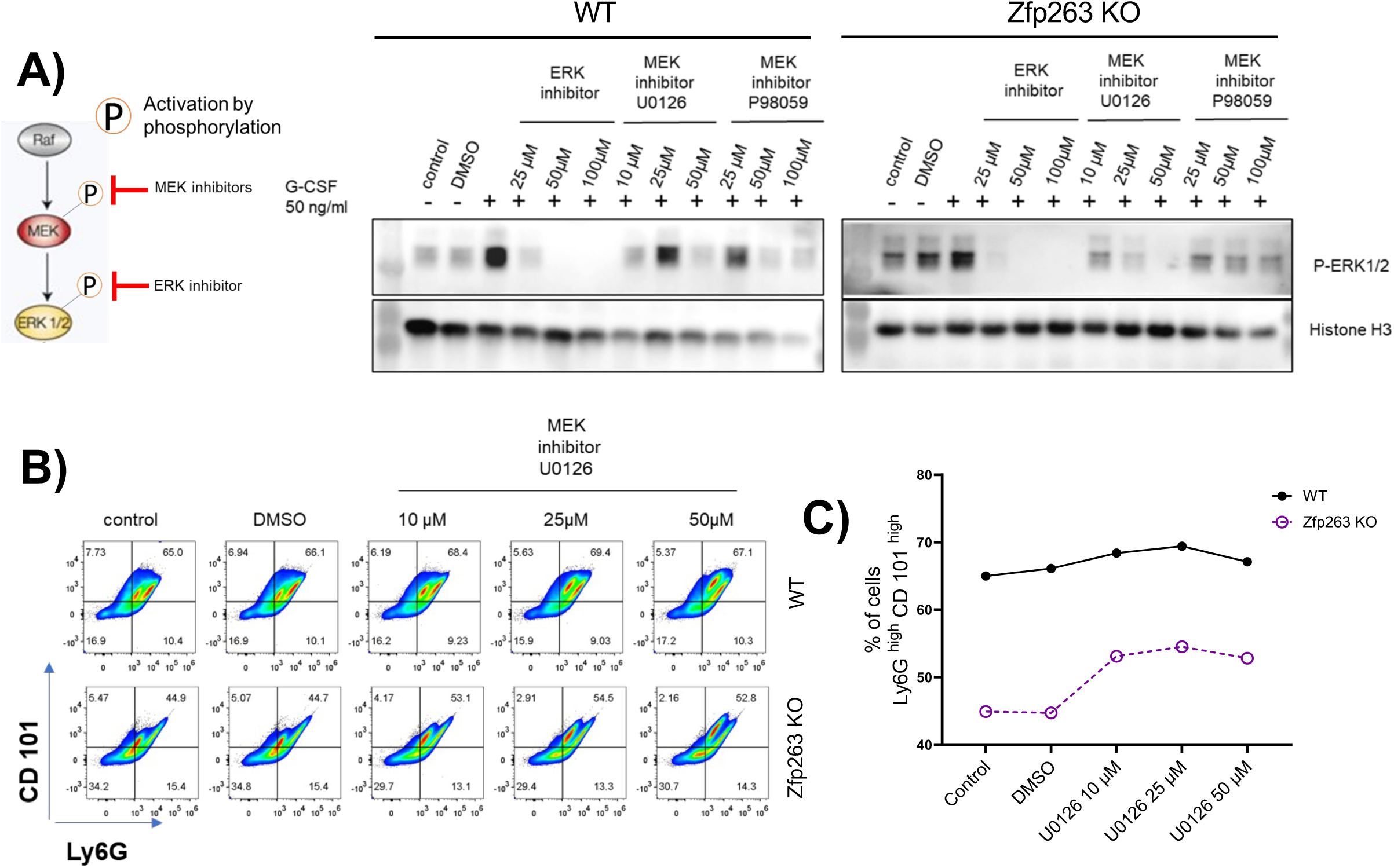
Late treatment of Zfp263KO neutrophils with ERK inhibitors partially restore maturation. A. Levels of phosphorylated ERK 1/2 were checked by western blot in HoxB8 progenitor cells (WT and Zfp263 KO) following 24 hr. treatment with ERK/MEK inhibitors at various concentrations. Histone H3 was used as a loading control for the experiment. Levels of ERK and p-ERK (normalized) were estimated by densitometry and plotted. B. Differentiation of HoxB8 cells (WT and Zfp263 KO) was checked on Day 5 after treating with different concentrations (10,25 and 50 µM) of the MEK inhibitor U0126 on Day 3 of differentiation. The expression of cell surface markers Ly6G and CD101 were estimated by flow-cytometry. Ly6G^high^ CD101^high^ cells were gated to determine the % of mature neutrophils. C. The % of Ly6G ^high^ CD101^high^ (mature neutrophils) was plotted for WT and Zfp263 KO cells at various concentrations of MEK inhibitor U0126 as shown in (B).

**Supplementary Figure S7:**
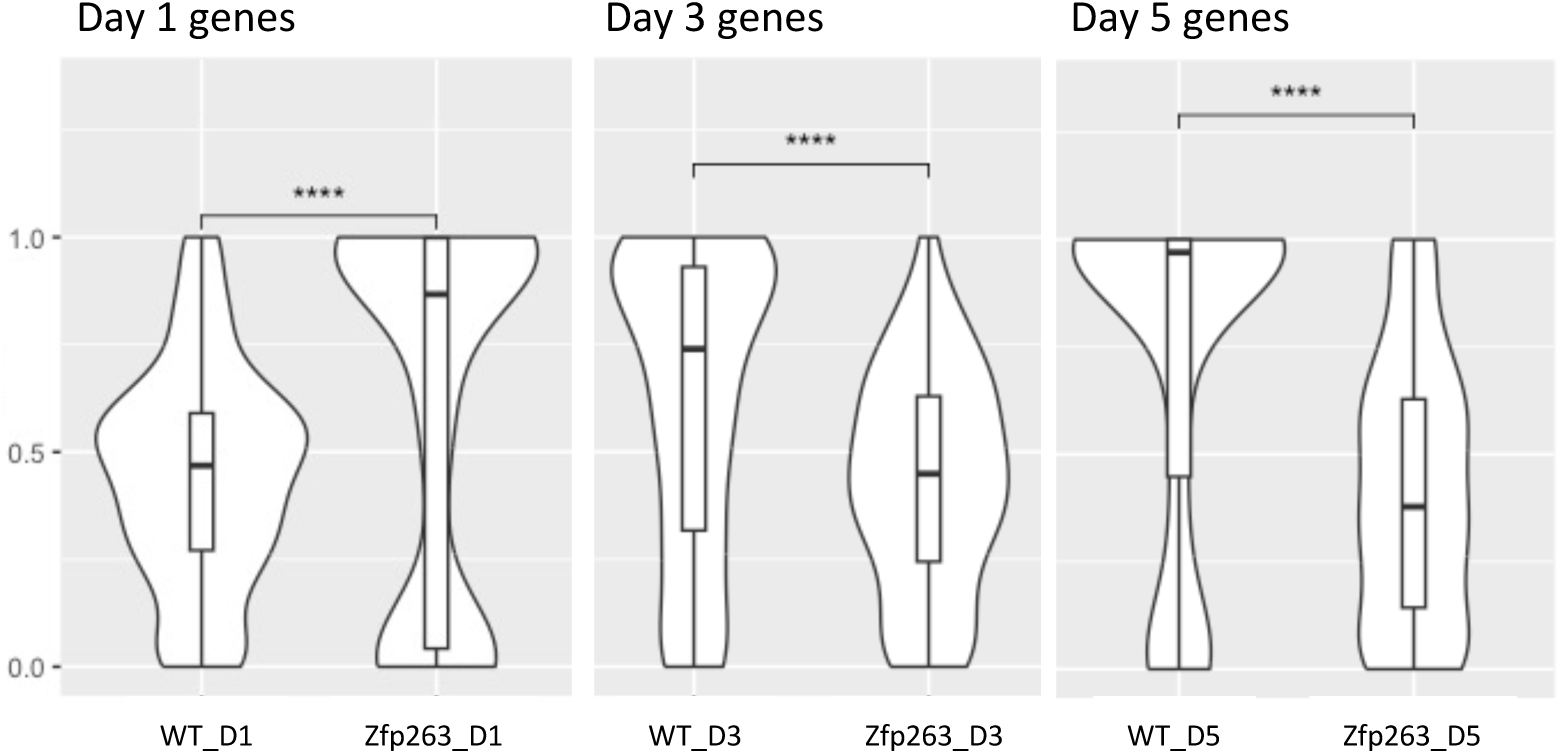
Zfp263 is a dual regulator (repressor and activator) of neutrophil gene expression. Gene expression trends of D1, D3, and D5 genes from the Zfp263 network (Figure 5A) in corresponding WT and Zfp263 KO cells

**Supplementary Figure S8:**
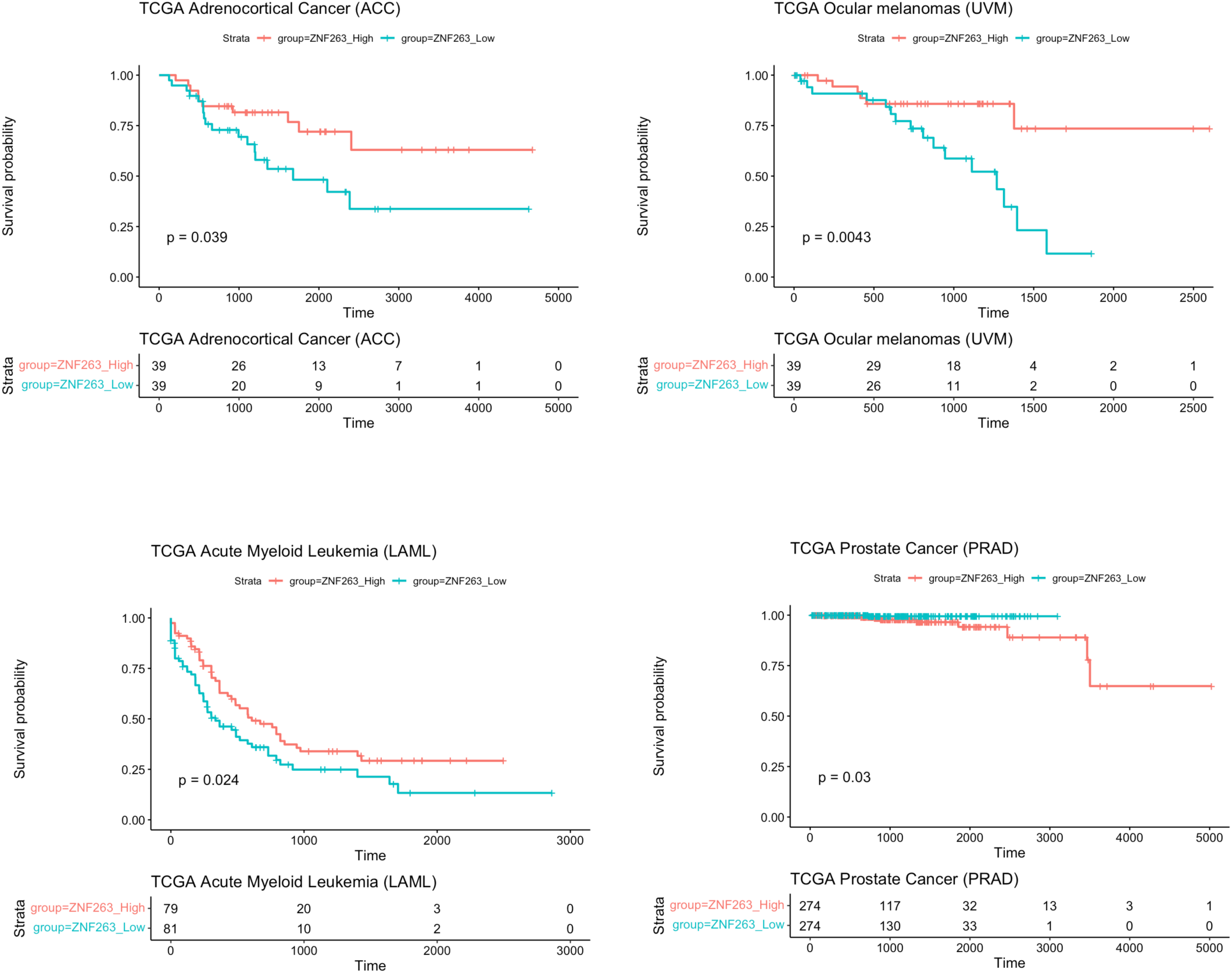
Survival analysis identifies cancer types positively and negatively correlated with ZNF263 expression levels. Survival analysis of various cancer types based on ZNF263 expression levels using Kaplan-Meier curves. The plots were generated using the “survfit” function from the ‘survival’ package in R, comparing the survival probabilities of patients with high and low expression of ZNF263 across different cancer types from the TCGA database.

**Supplementary method table 3.**

| constituent | concentration unit | concentration |
| --- | --- | --- |
| Hepes-KOH pH 7.5 | mM | 50 |
| NaCl | mM | 140 |
| EDTA | mM | 1 |
| glycerol | %(v/v) | 10 |
| NP-40 | %(v/v) | 0.5 |
| TritonX-100 | %(v/v) | 0.25 |

**Lysis Buffer 2**
| constituent | concentration unit | concentration |
| --- | --- | --- |
| Tris-HCl pH8.0 | mM | 10 |
| NaCl | mM | 200 |
| EDTA | mM | 1 |
| EGTA | mM | 0.5 |

**Lysis Buffer 3**
| constituent | concentration unit | concentration |
| --- | --- | --- |
| Tris-HCl pH8.0 | mM | 10 |
| NaCl | mM | 100 |
| EDTA | mM | 1 |
| EGTA | mM | 0.5 |
| Na-Deoxycholate | %(w/v) | 0.1 |
| Na-Lauroylsarcosine | %(w/v) | 0.5 |
| Triton-X 100 | %(w/v) | 1 |
| SDS | %(w/v) | 0.1 |

**low salt wash buffer**
| constituent | concentration unit | concentration |
| --- | --- | --- |
| Tris-HCl pH 8.0 | mM | 20 |
| NaCl | mM | 150 |
| EDTA | mM | 2 |
| TritonX-100 | %(v/v) | 1 |
| SDS | %(w/v) | 0.1 |

**high salt wash buffer**
| constituent | concentration unit | concentration |
| --- | --- | --- |
| Tris-HCl pH 8.0 | mM | 20 |
| NaCl | mM | 500 |
| EDTA | mM | 2 |
| TritonX-100 | %(v/v) | 1 |
| SDS | %(w/v) | 0.1 |

**LiCl wash buffer**
| constituent | concentration unit | concentration |
| --- | --- | --- |
| Tris-HCl pH 8.0 | mM | 10 |
| LiCl | mM | 250 |
| EDTA | mM | 1 |
| NP-40 | %(v/v) | 1 |
| Na-Deoxycholate | %(w/v) | 1 |

**Elution Buffer**
| constituent | concentration unit | concentration |
| --- | --- | --- |
| Tris-HCl pH 8.0 | mM | 10 |
| EDTA | mM | 1 |

**Supplementary method table 3.**
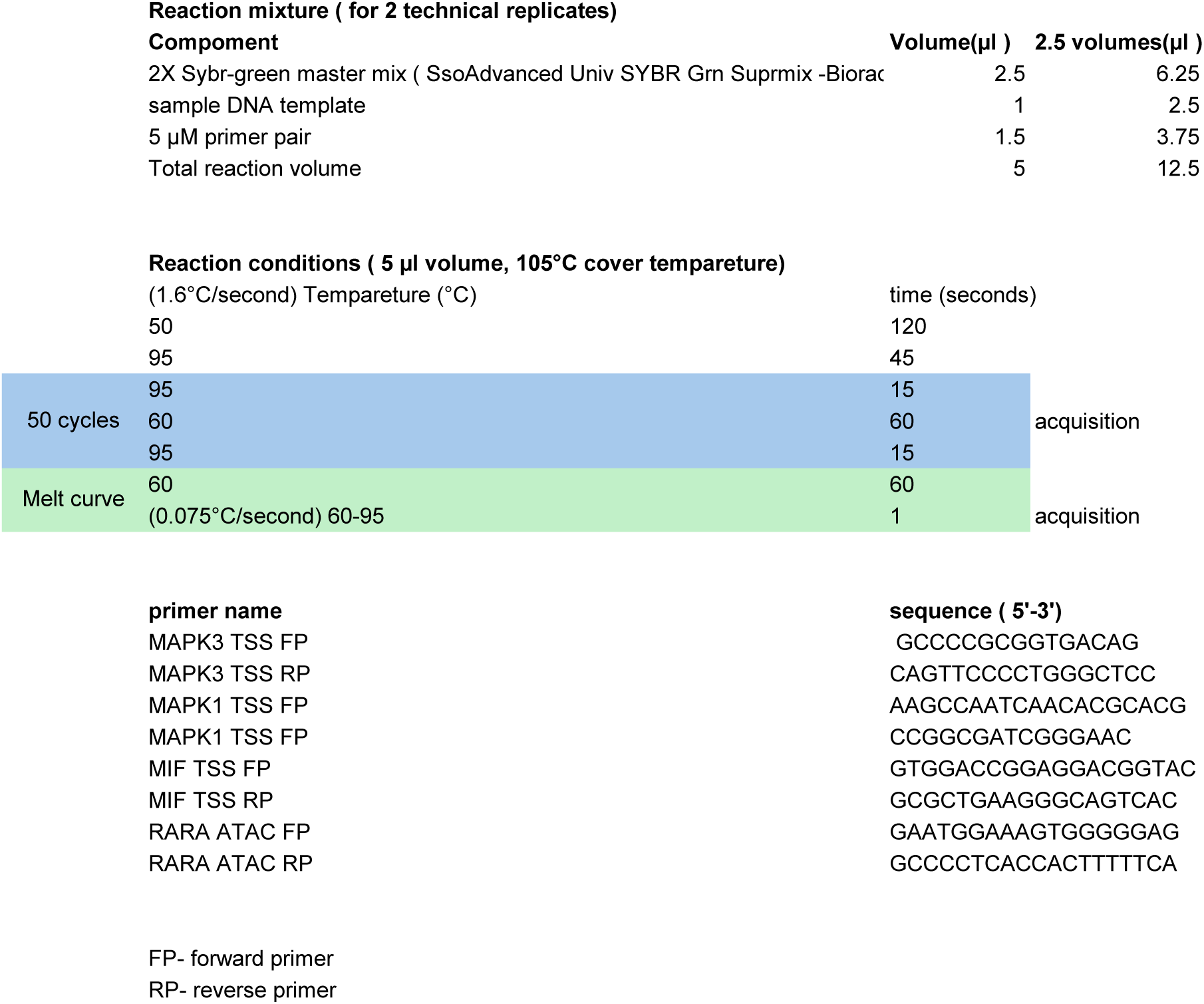

**Supplementary method table 3.**

| **Antibody target protein** | **Make and catalog number** | **Application** |
| --- | --- | --- |
| ZNF263 | SCBT-ZNF263 (8-YD33), Monoclonal Antibody, 100ug/ml | ChIP, Western Blot |
| IgG(Mock) | CST Mouse (G3A1) mAb IgG1 Isotype Control 5415 | ChIP |
| H3 | abcam-ab1791 | Western Blot |
| ERK1/2 | CST-P44/42 MAPK (ERK1/2) 137F5 4695S | Western Blot |
| phospho-ERK1/2 | CST-P-P44/42 MAPK T202/Y204 197G2 4377S | Western Blot |
| CD101 | Invitrogen-PE-Cy7 25-1011-80 | Flow cytometry |
| Ly6G | Biolegend- BV711 127643 | Flow cytometry |
| Ki67 | SCBT- Ki67 sc-23900-FITC | Flow cytometry |

